# Baseline growth rate shapes pollutant impacts in aquatic environments: Evidence from antibiotic responses in the harmful cyanobacterium *Microcystis*

**DOI:** 10.64898/2026.10.06.756710

**Authors:** Ganyu Feng, Zhipeng Duan, Changwei Hu, Juan Wang, Jie Xu, Yuejin Zhang, Xiang-Zhou Meng

## Abstract

Hormesis is an important consideration in chemical risk assessment across most aquatic environments. Evidence suggests that hormetic responses are shaped by organismal physiological processes. Accordingly, predictions of hormesis under environmentally realistic conditions should account for physiological traits. Yet the role of these traits remains largely unexplored. This study investigated how baseline growth rate modulates antibiotic responses in the harmful cyanobacterium *Microcystis* and the underlying mechanisms. Results showed that at an antibiotic concentration of 200 ng L^-1^ (representing environmental concentration levels), slow-growing *Microcystis* populations exhibited an obvious growth enhancement more than 4-fold greater than that of fast-growing populations, and this enhancement was positively correlated with cellular reserves. A baseline growth rate below 0.19 day^-1^ emerged as the threshold below which hormetic responses were consistently observed. This study provides pioneering insights into how organisms’ physiological states influence pollutant-induced hormetic responses in aquatic environments. Our findings point to a central principle for extrapolation that laboratory-derived toxicological responses may not translate directly to the field unless key physiological traits of the organisms are explicitly considered. In this regard, baseline growth rate and energy reserves may be key physiological traits for reliably predicting toxic effects at the population level. This study also makes an initial attempt to conceptualize a physiology-anchored framework for strengthening risk assessment of low-level emerging contaminants.

## 1. Introduction

The impacts of emerging contaminants in the water environment are gradually being brought into focus (Ruan et al., 2023). One of the cornerstones is the hormesis theory that the most fundamental dose-response curve in the real word is not linear or threshold-based, but U-shaped (Agathokleous, 2025). Theoretically, incorporating hormesis into ecological risk assessment (ERA) is feasible (Rillig et al., 2025; Sun and Wu, 2026). However, field conditions often diverge from laboratory predictions, which is a gap that may stem from organism’s growth states (Tang et al., 2022). Most responses incur biological costs (Schavemaker and Lynch, 2025), particularly the depletion of cellular resources (Jager et al., 2013). Conversely, organisms in good growth condition may exhibit hormesis more fully (Tang et al., 2022). However, the connection between physiological traits and hormesis remains largely unexplored.

Cyanobacterium *Microcystis*, notable for its notorious role in causing harmful water blooms, is widely used as a model organism in hormesis-related researches (Geng et al., 2025). Evidence indicates that low levels of chemicals can stimulate its growth and enhance toxin production (Agathokleous et al., 2022). Notably, antibiotics can trigger hormesis effect at concentrations as low as several ng L^-1^, while other chemicals usually require μg L^-1^ or even mg L^-1^ to produce similar effects (Agathokleous et al., 2022; Ve Van et al., 2022; Wang and Tao, 2025). This distinction positions antibiotics as a particularly notable class of pollutants, capable of exerting effects at concentrations commonly found in the water environment. Here, *Microcystis* and antibiotics constitute an ideal test pair, and the findings are highly relevant to real-world aquatic environments. Specifically, these findings can reveal how key physiological traits affect the hormesis effect.

In an earlier work on this organism-chemical pair, a lower baseline growth rate (defined as the pre-hormesis growth rate) was observed to correlated with substantially stronger hormesis effects (see Supplementary **Fig. S1**a,b). This observation raises the hypothesis that baseline growth rate may underlie the observed variability in dose-responses. At present, however, there is a scarcity of reliable studies that elucidate the regulatory factors potentially influencing the hormesis effect. A major barrier to cumulative progress lies in the poor comparability across existing studies, which vary widely in strains, culture media, test chemicals, dosage regimens, and exposure windows (Donzelli et al., 2025). Here, a plausible explanation is that enhanced growth comes at the expense of other metabolic processes (Zhu and Dai, 2024). Rapid growth entails increased cell turnover, accelerated loss of carbon (C) and nutrients, and reduced investment in storage reserves (Huang et al., 2019). Such a trade-off eventually may hit a biological ceiling, limiting further response to stressors (Moreno-Paz et al., 2022). Building on this reasoning, our study first examined the relationship between hormetic responses and baseline growth rate, then interpreted it in terms of energy reserves.

Stoichiometric perspective centered on carbon, nitrogen (N), and phosphorus (P) is fundamentally viable because stoichiometry is consistent with energy reserve budget framework (Sperfeld et al., 2017) and is linked to functional responses (Moody et al., 2025). For most phytoplankton, the three primary biogenic elements constitute approximately half of their dry mass (Maberly et al., 2021). Growth is fueled by the portion of the cellular element quota that surpasses the minimum requirement (Zonneveld, 1996; Klausmeier et al., 2004). External factors often fail to stimulate growth once key elements near their minimum cellular quotas (Verspagen et al., 2014), analogous to the Liebig’s law of the minimum (Klausmeier et al., 2004). The potential roles of cellular reserves linked to different elements have received relatively limited attention in previous studies, and exploring this dimension represents a promising avenue for advancing our understanding.

This study aimed to clarify the hormesis effects associated with baseline growth rates and the role of underlying reserve constraints. Specifically, we investigated *Microcystis* growth under hormesis induced by low-level antibiotics (sulfamethoxazole, tetracycline, roxithromycin, and ofloxacin) across varied baseline growth rates. Our results demonstrate this relationship and highlight the key factors in enabling hormesis. The findings offer a forward-looking view on assessing chemical impacts in the water environment, particularly for improving the scientific basis of evaluating pollution responses *in situ*. Our study also reveals a previously overlooked determinant of hormesis variation. This determinant can be translated into a practical tool for refining ERA and management approaches for water pollutants.

## 2. Materials and methods

### 2.1. Semi-continuous cultures and control of baseline growth rate

To maintain relatively stable baseline growth status, we employed semi-continuous cultivation with daily dilution in standard BG11 medium (Supplementary **Fig. S1**c,d,e), a method that can be conceptualized as a series of short-cycle batch experiments (LaRoche et al., 2010). The dilution rate (DR) ranged from 0.05 to 0.20 day^-1^. *Microcystis* sp. strain PCC 7806 (Pasteur Culture Collection, Paris, France) was selected as the test organism. Before inoculating, 1-L Erlenmeyer flasks were scrubbed three times with methanol, acetone, and dichloromethane, followed by drying. They were then washed with ultrapure water and dried again. Each flask was sealed with high-temperature resistant tissue culture sealing film (Bkmam, Changde, China), which allowed gas exchange. The flasks were then autoclaved at 121°C for 30 min. Stocks of culture media (standard BG11) were sterilized by autoclaving, kept at 4°C in the dark, and heated to 25°C before use. Each flask contained 300 mL of BG11 medium and was located in an illuminating incubator (Huitai QHQ-250, Shanghai, China) set at 25°C under a 12-hour day/night cycle, with a constant incident light intensity of 50 μmol photons m^-2^ s^-1^ and a relative air humidity of 60% to 70%. Daily dilutions were carried out at the end of the dark period by taking samples from well-mixed cultures and supplementing with an equal volume of fresh medium. Samples taken were used for the relevant measurements. Flasks were shaken daily and their positions were exchanged with each other.

### 2.2. Exposure experiments

The cell concentration of each semi-continuous culture was maintained at a stable growth state before any further exposures (Fig. 1a,b). Then, specific doses of quaternary antibiotic mixtures (QAMs) were added. The QAMs included sulfamethoxazole, tetracycline, roxithromycin, and ofloxacin. All four antibiotic categories are frequently detected in surface water (Lyu et al., 2020). Axenic stock solutions of QAMs were prepared following the methodology described in a previous study (Feng et al., 2023), using powders purchased from Macklin Co., Ltd. (Shanghai, China) with a purity exceeding 98%. Additional stock solutions were added to the cultures daily to compensate for the losses due to dilution and degradation (Liu et al., 2012). Cultivation conditions were kept consistent with those described in the semi-continuous cultures. Various dilution rates, doses, and nutrient conditions were selected for exposure experiments. In the two modified media, the levels of N and P were reduced to 10% of those in the standard BG11 medium, and the concentrations of sodium and potassium were balanced by the addition of NaCl and KCl, respectively. Each exposure incubation was prepared with at least two biological replicates.

**Fig. 1.**
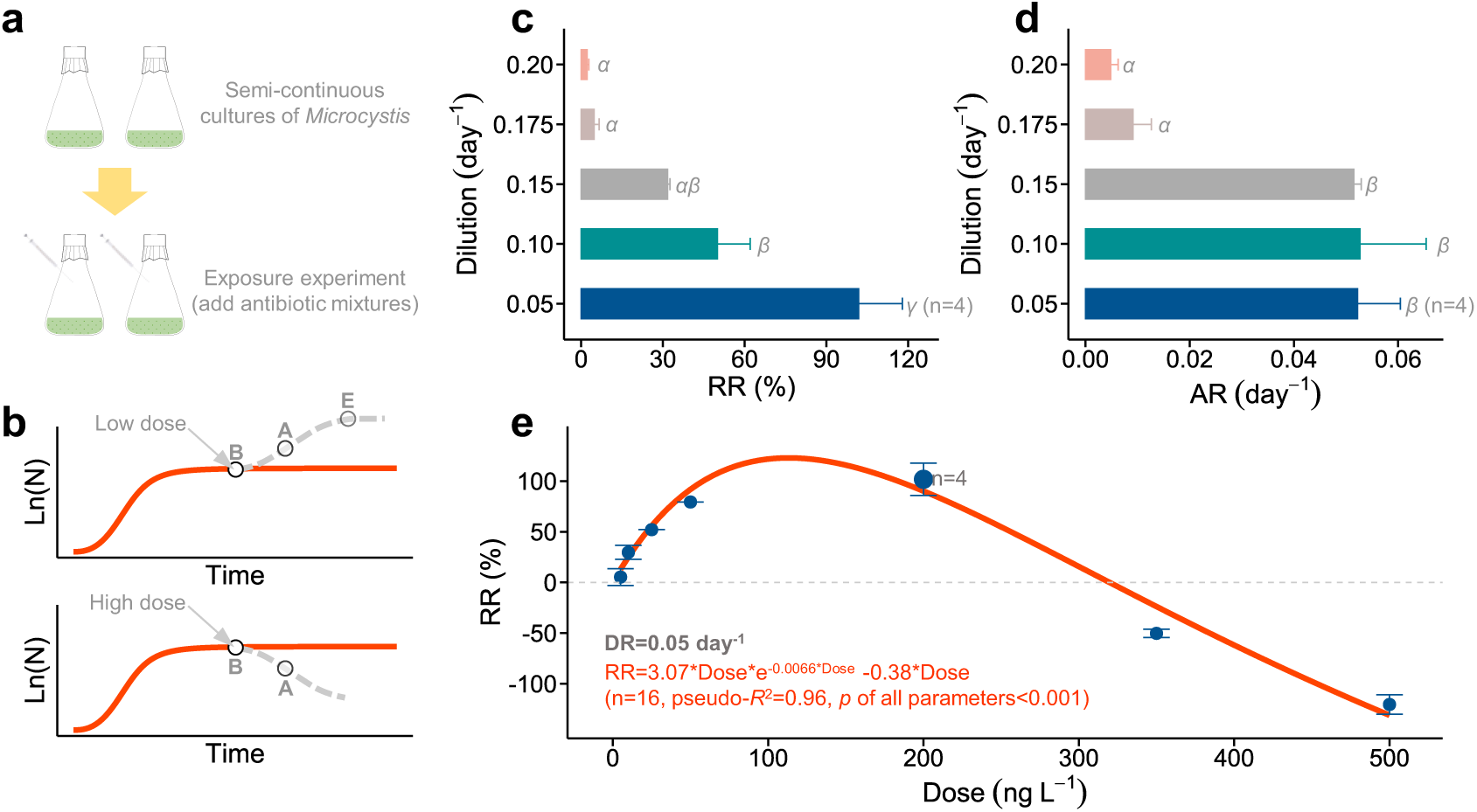
Responses of growth after exposures. **a** Schematic diagram of the experimental process for semi-continuous cultures and exposures. **b** Expected changes in cell concentration under low and high doses (grey dashed curves, respectively), compared to the unexposed control (solid red curve). Labels ‘B’, ‘A’, and ‘E’ denote time points before, during, and after dose exposure, respectively. **c** Relative response (RR) in growth rate during exposure to 200 ng L^-1^, calculated using Eq. (S4) across different dilution rates (DRs). Effective hormetic response is set at RR>10% (Agathokleous, 2025). The administered dose (200 ng L^-1^) is environmentally relevant (Lyu et al., 2020). **d** Absolute response (AR) in growth rate during exposure to 200 ng L^-1^, calculated using Eq. (S5) across different DRs. Greek letters (*α*, *β*, and *γ*) indicate a significant difference (*p* < 0.05) between dilution rates. **e** RR under different antibiotic doses. Exposures under a dilution rate of 0.05 day^-1^ and a dose of 200 ng L^-1^ were repeated twice, which doubled the sample size (i.e., n=4); for other exposure experiments, n=2. The growth curves before and after exposures are shown in Supplementary **Fig. S2**a,b.

A stock solution of solid phase extraction (SPE) from Lake Taihu replaced QAMs in the exposures (Supplementary **Text S1**). The cultures were divided into test tubes (2.5 mL per tube) for further exposures. We set up four experimental groups: STD (standard BG11 medium), STD+SPE, -N (N-limited medium), and -N+SPE, with ten replicates per group. The designation ‘+SPE’ indicates that SPE stock was added to the medium. For the +SPE groups, 50 μL of SPE stock was added to each test tube and mixed well. For the two control groups (STD and -N), 50 μL of sterilized ultrapure water was added to each test tube instead. Each group was prepared with ten biological replicates. The incident light intensity was 25 μmol photons m^-2^ s^-1^ and other cultivation conditions were consistent with those of exposure experiments. Optical density at a wavelength of 680 nm (measured using a Spectrophotometer Puxi T6, Beijing, China) in each test tube was recorded 48 hours after exposure onset.

### 2.3. Measurements

After the culture was injected into a hemocytometer and allowed to settle completely, the cell concentration was enumerated under a microscope (Olympus, Tokyo, Japan) at a magnification of 200 ×. Each pair of connected cells in the division phase or after division completion is counted as two cells. The absolute response AR_j_ and relative response RR_j_ in growth after ‘j’ hours of exposure onset were calcul ated using the following formulas:

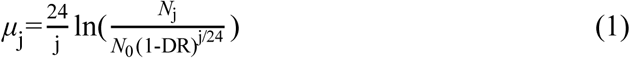

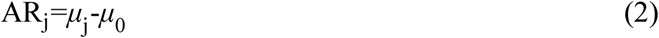

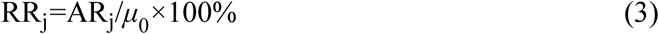

where, *N*_j_ and *μ*_j_ are the cell concentration and average specific growth rate after ‘j’ hours of exposure onset (e.g., 24 h, 48 h, 72 h …), respectively. *N*_0_ represents the baseline cell concentration before exposure. *μ*_0_ is the baseline growth rate, which depends on the dilution rate (refer to Supplementary **Fig. S1**c). This study compared differences at specific points in time (e.g., j = 48), unless otherwise stated (calculations for the entire duration of exposure are listed in Supplementary **Text S2)**.

Samples for the analysis of cellular C, N, and P were collected on clean 0.45-μm glass fiber filters. The volume of samples ranged from 10 to 40 mL, depending on the dilution rates. The filters were then dried for 24 h at 70°C and stored in an amber desiccator. For the analysis of C and N, a portion of each filter was taken using a clean hole puncher and analyzed using a UNICUBE organic elemental analyzer (Elementar Analysensysteme GmbH, Langenselbold, Germany). P was analyzed using the ammonium molybdate spectrophotometric method (GB 11893-89, China) after combusting a portion of each glass fiber filter at 400°C for 120 min. The cellular content of C, N, and P was determined by dividing the total element content of the sample by the number of cells. Variations in cellular content of element *Q* during specific length of time (from time 0 to time j) was calculated using the following formula:

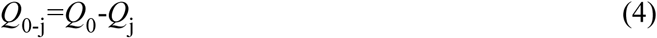

In addition, the concentrations of residual dissolved N and P in the filtrates of cultures were measured according to the methods HJ 636-2012 (Alkaline potassium persulfate digestion UV spectrophotometric method, China) and GB 11893-89, respectively. IC in the filtrates was measured using a total organic carbon analyzer (Jena Multi N/C 3100, Freistaat Thüringen, Germany). Cell diameters were determined with ImageJ (v. 1.54f) under 400× magnification. Transcriptomes from two exposure experiments, conducted with a dose of 200 ng L^-1^ and different dilution rates of 0.05 and 0.20 day^-1^, were sampled and then analyzed (Supplementary **Text S3**). Raw RNAseq reads for differential gene expression analyses have been submitted to NCBI’s SRA database (http://www.ncbi.nlm.nih.gov) under BioProject PRJNA1153622.

### 2.4. Statistical analyses

Statistical analysis was conducted using R 4.2.3 (https://www.r-project.org/). The data were presented as mean ± standard deviation, unless otherwise stated. Differences between treatments were analyzed using one-way analysis of variance (ANOVA) with Tukey’s HSD test. The significance level was set at *p* < 0.05. PCA was performed using the ‘PCA’ function in the ‘FactoMineR’ package. Curve fitting was conducted using the ‘nls’ function in the ‘stats’ package. Heatmap analysis was performed using the ‘pheatmap’ package. Venn analysis was performed using the ‘ggvenn’ package and visualized using the ‘eulerr’ package. The statistical analysis of transcriptomes is described in Supplementary **Text S3**.

## 3. Results

### 3.1. Key physiological trait: Baseline growth rate shapes pollutant impacts

The baseline growth rate at DR = 0.05 day^-1^ approximated the reported *in-situ* rate (Browning and Moore, 2023). A pronounced hormesis effect was evident at lower DRs, whereas cultures at high DRs (0.175 and 0.20 day^-1^) displayed minimal sensitivity (see Fig. 1c,d; Supplementary **Fig. S2**a). Compared with fast-growing populations, slow-growing *Microcystis* exhibited >4-fold and >5-fold greater growth enhancement as measured by the AR and RR indices, respectively (Fig. 1c,d). It is noteworthy that a marked decrease in both absolute and relative response occurred at a DR of 0.175 day^-1^ (Fig. 1c,d). So far, we have preliminarily validated the phenomenon that an excessively high baseline growth rate leads to the disappearance of hormesis effect. Additionally, owing to its environmental relevance and the clear sensitivity exhibited, the condition of DR = 0.05 day^-1^ was investigated for dose-response effects, revealing an increase in growth rate under low antibiotic exposure but a decline at higher doses (Fig. 1e).

### 3.2. Cellular reserve availability underpins the hormetic response

We investigated the responses of biomass (measured as cellular dry weight) and key stoichiometric traits to exposure at 200 ng L^-1^ under conditions without strict nutrient limitation. Cellular dry weight, carbon and nitrogen content, and cell diameter all decreased with longer exposure time, particularly at low dilution rates (Supplementary **Fig. S3**a-c,h-k). Hormetic responses showed positive correlations with reductions in carbon (adj-*R*^2^ = 0.78, *p* < 0.001) and nitrogen content (adj-*R*^2^ = 0.62, *p* < 0.01; Fig. 2a-d and Supplementary **Table S1**). We further observed that this trend is not confined to growth enhancement triggered by hormesis; instead, it generally occurs under conditions of accelerated growth (Fig. 2e,f and Supplementary **Fig. S4**). Moreover, carbon and nitrogen quotas (per cell volume) appeared to exceed the levels required for growth at low growth rates (Supplementary **Fig. S3**e,f,g). These results suggest that the hormesis effect in *Microcystis* is more strongly linked to carbon and nitrogen quotas (Supplementary **Table S1**).

**Fig. 2.**
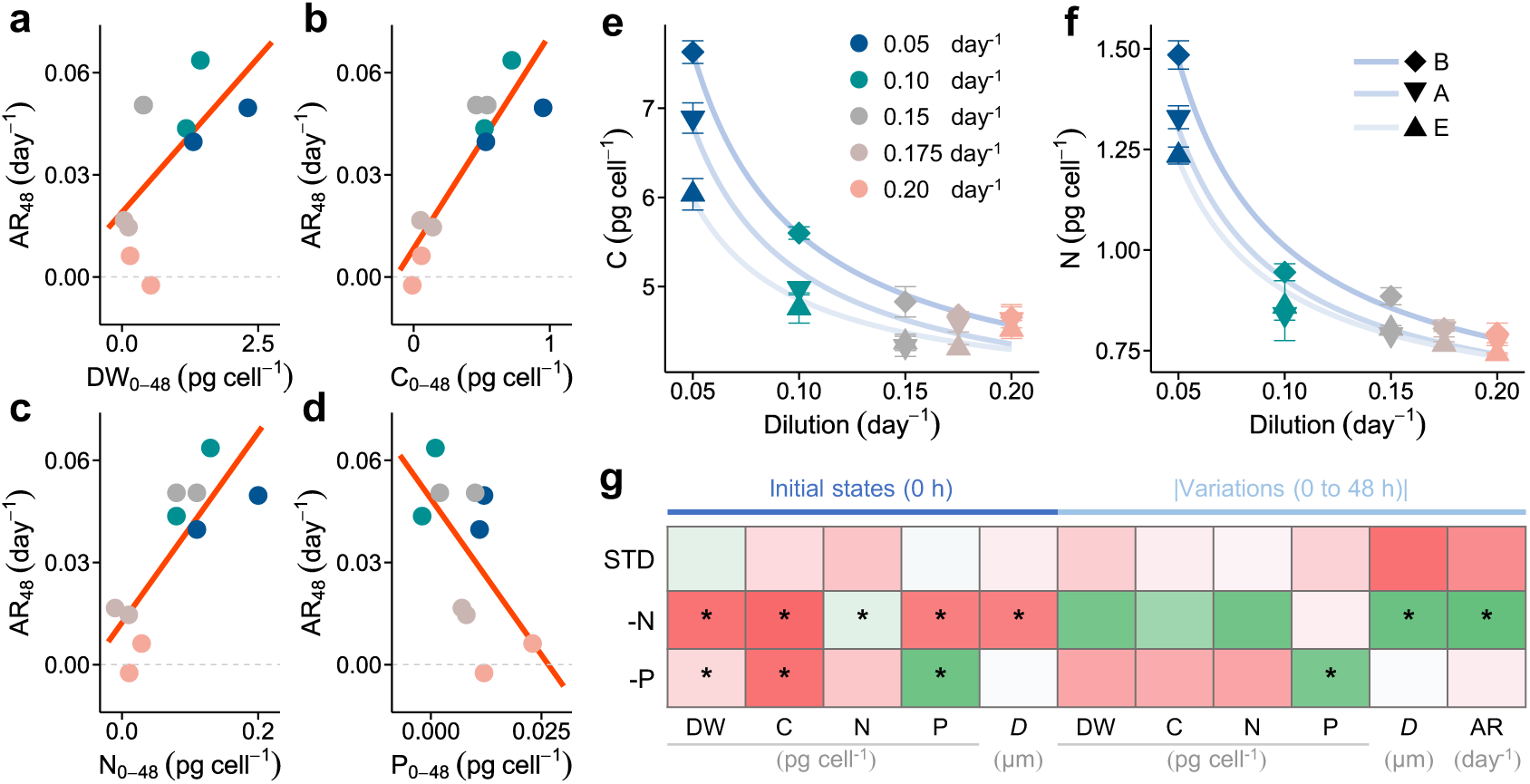
Relations between growth response and cellular reserves. **a** Relationship between absolute growth rate response after 48 hours (AR_48_) and reduction in cellular dry weight (DW_0-48_) under exposure to 200 ng L^-1^. **b** Relationship to reduction in cellular content of carbon (C_0-48_). **c** Relationship to reduction in cellular content of nitrogen (N_0-48_). **d** Relationship to reduction in cellular content of phosphorus (P_0-48_). Data points represent individual biological samples rather than mean values. Details on the linear fitting results are listed in Supplementary **Table S1**. **e** and **f** Cellular contents of carbon and nitrogen under different dilution rates. ‘B’ for pre-exposure, ‘A’ for post-exposure, and ‘E’ for end of the experiment. Similar to the growth curves, the cellular characteristics are also assumed to be constant at node ‘B’ and changes at ‘A’. The details of the curve fitting are presented in Supplementary **Fig. S4**. **g** Cellular initial states and absolute values of variation among groups with three nutrient supply conditions (STD, -N, and -P). The heatmap provides a comparison among groups at the level of each indicator. The color was determined based on the mean value of the biological duplicates and was assigned using Excel (Microsoft Office 2016). In each column, the warmer the color, the larger the value. Asterisk (*) indicate a significant difference (*p* < 0.05) to STD. ‘*D*’ represents cell diameter. The growth curves of the STD, -N, and -P groups are shown in **Fig. S2**a and **Fig. S5**a.

We further modified BG11 medium to generate nitrogen-limited and phosphorus-limited treatments. The final residual concentrations of all nitrogen and phosphorus are detailed in Supplementary **Table S2**. All cultures were maintained at a dilution rate of 0.05 day^-1^ and exposed to a dose of 200 ng L^-1^. After dosing, compared to the STD group, the -N group maintained constant reserves and lost the ability to mount a hormetic response (Fig. 2g; Supplementary **Fig. S5**). In contrast, the -P group still exhibited growth-promoting effects (with an AR_48_ of 0.03 day^-1^) along with alterations in dry weight, cellular nitrogen content, and cell diameter (Supplementary **Fig. S5**). These observations show that although limiting the cellular quota of either nitrogen or phosphorus suppresses hormesis, the effect of nitrogen limitation is stronger, especially over short timescales. Furthermore, eliminating light energy input also abolished the hormetic response (Supplementary **Fig. S6**), further confirming a strict dependence on external resource availability. Collectively, these results reveal that hormesis is abolished under severe growth limitation, whereas in its absence, the magnitude of hormesis scales with the cellular potential for further accelerated growth (indexed by intracellular carbon and nitrogen content) such that a greater potential elicits a stronger hormetic effect.

### 3.3. Transcriptomic features of growth acceleration

Under nutrient-sufficient culture conditions, gene expression displayed similar patterns in response to 200-ng L^-1^ antibiotic exposure (A vs. B) and to an increased dilution rate (0.20 day^-1^ vs. 0.05 day^-1^; Fig. 3a,b,c and Supplementary **Fig. S7**). Most differentially expressed genes (DEGs) were upregulated in 0.05:A/B and B:0.20/0.05 comparisons, but downregulated in 0.20:A/B (Supplementary **Fig. S8**). Approximately half of the DEGs were shared between the 0.05:A/B and B:0.20/0.05 comparisons (Fig. 3d and Supplementary **Table S3**), and these overlapping genes exhibited an approximately 1:1 linear relationship (Fig. 3e). In contrast, at high dilution rates, antibiotic-induced changes in DEG levels (see 0.20:A/B) correlated poorly with other antibiotic or dilution conditions (Supplementary **Fig. S9**a,b). These patterns suggest a transcriptomic convergence between the hormesis effects of antibiotics and those of growth acceleration induced by an increased dilution rate.

**Fig. 3.**
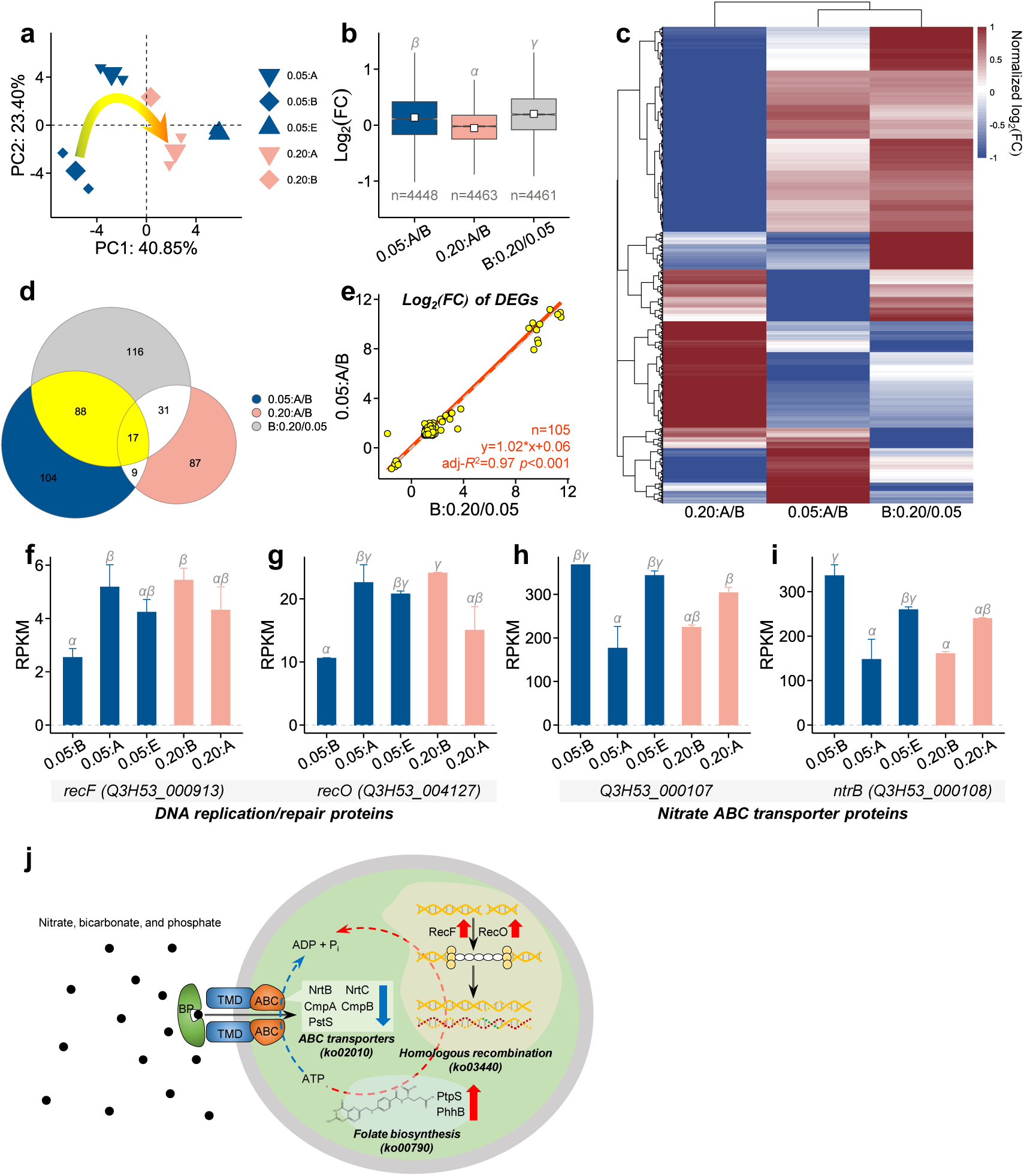
Comparison of transcriptomes between antibiotic-stimulated growth and growth under various dilutions. **a** PCA plot of variations in the RPKM of all mapped genes. Samples are labeled in the format ‘dilution rate (0.05 or 0.20) : time point (B, before; A, after; E, end of exposure)’. **b** FC of all mapped genes among comparisons. For example, ‘0.05:A/B’ is the 0.05:A vs. 0.05:B comparison aimed to find variations in transcriptomes before and after exposure under a dilution rate of 0.05 day^-1^ (i.e., exposure-stimulated growth). ‘B:0.20/0.05’ is the 0.20:B vs. 0.05:B comparison focused on growth driven by dilution. **c** Heatmaps for all mapped genes. **d** Venn plot of DEGs. A total of 252, 218, and 144 DEGs were identified in the B:0.20/0.05, 0.05:A/B, and 0.20:A/B comparisons, respectively. The intersection between the first two comparisons (yellow) contains the common set of DEGs. **e** Scatter plots to compare exposure-stimulated growth and growth driven by dilution based on the common set of DEGs in panel **d**. The solid line represents the best fit for all data points in the panel, along with the equation of the fitted function. The slope of the dashed line is 1 and the intercept is 0. The details of the 105 DEGs are listed in Supplementary **Table S3**. **f** and **g** Gene expressions of DNA replication/repair proteins in enriched KEGG pathway ko03440. **h** and **i** Gene expressions of nitrate ABC transporter proteins in enriched KEGG pathways ko02010 and ko00910. The details of the enriched KEGG pathways are listed in Supplementary **Table S4**. Greek letters (α, β, and γ) indicate a significant difference (*p* < 0.05). **j** the transcriptomic changes in key metabolic and regulatory mechanisms of *Microcystis* under antibiotic stress. It highlights three core biological processes (transmembrane transport, DNA damage repair, and folate biosynthesis), with red arrows indicating the upregulation of related gene expressions.

During hormesis-induced growth acceleration, upregulated DEGs were enriched for homologous recombination and folate biosynthesis, while downregulated DEGs were enriched for nitrate ABC transporters and nitrogen metabolism (**Table S4**). Increasing the dilution rate alone also resulted in enrichment of DEGs in homologous recombination and nitrogen metabolism pathways. Under high baseline growth rate conditions, however, antibiotic exposure led to a different transcriptomic profile in which upregulated DEGs were enriched in quorum sensing and downregulated DEGs in sulfur metabolism. Notably, expression of DNA replication/repair genes (*recF* and *recO*) increased markedly after exposure at the low dilution rate but remained stable at the high rate (Fig. 3f,g). A similar trend was observed for folate synthesis genes such as Q3H53_002006 (Supplementary **Fig. S10**a). By the end of exposure period, *recF* and *recO* expression remained elevated, while the folate-related genes returned to baseline. Expression of ABC transporter genes, including *ntrB*, Q3H53_000107, and Q3H53_003417, decreased rapidly under low dilution exposure but stayed relatively constant at the high rate (Fig. 3h,i and Supplementary **Fig. S10**e-h). More photosynthetic genes were also downregulated during early exposure or at high growth rates (Supplementary **Fig. S11**).

### 3.4. Toward in situ extrapolation: Responses to field-collected lake water

We further tested the response to a mixture of organic components from Lake Taihu, China’s third largest freshwater lake that is frequently plagued by cyanobacterial blooms, and contaminated with trace levels of various pollutants such as antibiotics (Li et al., 2020). The growth rate increased significantly in the STD+SPE group compared to the STD group, by 37.2%±4.1% (**Fig. S12**). In contrast, growth variation was negligible in nutrient-limited groups (-N+SPE vs. -N), with only a -0.4%±7.7% change. These results suggest that Lake Taihu water can stimulate *Microcystis* growth when the cells are physiologically normal, have a low baseline growth rate, and possess adequate energy reserves.

## 4. Discussion

Understanding how physiological states modulate hormesis is central for assessing the impacts of emerging contaminants. This study investigated a typical *Microcystis*-antibiotic research pair, where aquatic relevant concentrations trigger hormetic responses (Agathokleous et al., 2022; Ve Van et al., 2022). Slow-growing populations (DR = 0.05 – 0.15 day^-1^) exhibited substantially >4-fold greater growth stimulation than faster-growing ones (Fig. 1c,d). A baseline growth rate below 0.19 day^-1^ (i.e., DR < 0.175 day^-1^) represents a threshold worthy of further investigation for determining the likelihood of positive responses. Our results establish a significant negative association between the extent of hormesis and the baseline growth rate, pointing to a greater hormetic susceptibility in slower-growing populations.

### 4.1. Possible mechanisms behind the relationship between baseline growth rate and hormesis effect

The transcriptomic response suggests that hormesis-associated changes partly overlap with a general response to accelerated growth. Under nutrient-sufficient conditions, antibiotic exposure at the low dilution rate and an increased dilution rate shared approximately half of their differentially expressed genes, with the shared expression changes following an approximately 1:1 relationship (Fig. 3a-e). This overlap was weak under the high-growth-rate condition, where the antibiotic response differed from the other comparisons (**Supplementary Fig. S9**a,b). Consistent with this pattern, cellular C and N levels declined not only during hormesis but also under other accelerated-growth conditions (Fig. 2e,f). These results suggest that low-rate cells may hold C and N reserves that can support faster growth (Droop, 1973), but that the growth response is accompanied by their depletion.

Generally, hormesis stimulates metabolic activity and upregulates ATP synthase, thereby boosting reserve consumption (Wang and Tao, 2025), and our results highlight DEGs linked to growth and reserves. Although hormesis accelerates growth and reserve consumption, it does not enhance assimilation capacity (Wang et al., 2023; You et al., 2023). ABC transporter genes including ntrB, Q3H53_000107, and Q3H53_003417 decreased rapidly under low dilution exposure but stayed relatively constant at the high rate (Fig. 3h,i), while more photosynthetic genes were downregulated during early exposure or at high growth rates (Supplementary **Fig. S11**). These transcriptomic features indicate that *Microcystis* downregulated other cellular metabolic activities, such as nitrogen-related transport (Abdullah et al., 2025), and increased investment in maintaining the accuracy of genetic information when growth rates were enhanced, supporting the idea that trade-offs between different traits can prevent simultaneous hormetic improvement in all traits, especially in organisms with weak homeostasis (Sousa et al., 2010; Erofeeva, 2024). Therefore, in the absence of strict growth limitation, the shift in gene expression priority from resource acquisition to cell division underlies the observed trade-off between growth promotion and the drawdown of cellular reserves.

Here, Fig. 3j provides a possible transcriptomic basis for this trade-off: (i) greater investment in RecFOR-mediated homologous recombination and DNA repair (Rajaram et al., 2020); (ii) the increased expression of a folate-biosynthesis gene may also support folate-dependent purine and thymidylate synthesis, thereby supplying nucleotides for DNA synthesis during repair and cell proliferation (Handa et al., 2009; Mills et al., 2020); and (iii) nitrate-transport and nitrogen-metabolism genes were downregulated, similar to changing nutrient conditions (Harke and Gobler, 2013). These patterns are consistent with a shift in expression priority from resource acquisition toward DNA maintenance and division. Such reallocation could raise population growth rate while reducing cellular resource quotas and cell size, as growth rate, nutrient supply, cell size, and elemental quotas are interrelated in cyanobacteria and phytoplankton (Garcia et al., 2016; Inomura et al., 2020).

### 4.2. Controlling elements of hormetic potential

Bioelements play a critical role in characterizing reserve dynamics during the hormetic response, as demonstrated by our results (**Fig. 2**). In this process, carbon-rich energy reserves are essential. Nitrogen, as a central component of metabolism, influences the efficiency of reserve mobilization. The phosphorus content of *Microcystis* cells showed a good correlation with growth rate (consistent with the Droop model), while carbon and nitrogen exhibited greater variability and tended to accumulate at low growth rates, forming unallocated reserves (**Fig. S3**e-g). While phosphorus is generally considered the ultimate long-term limiting factor, short-term growth is frequently more sensitive to the availability of carbon and nitrogen (Seelen et al., 2025). From a cellular budget perspective, carbon and nitrogen are closely tied to energy mobilization, and may therefore govern the immediate response upon exposure. Additionally, cellular elemental ratios exceeding the thresholds of C_125_:P_1_ (molar) and the Sterner ratio (N_20_:P_1_) may serve as potential indicators of hormesis capability (**Fig. S3**l-n). It should be noted that nutrient deprivation markedly inhibited or attenuated the hormetic response (Fig. 2g). Strict nitrogen stress can induce the degradation of pigments, promote the accumulation of carbohydrates and lipids, and lead to the production of inhibitory metabolites, ultimately driving cells into a dormant state (Tandeau de Marsac and Houmard, 1993). The absence of hormesis under -N conditions may be partly attributable to insufficient activation of key enzymes, such as glucose-6-phosphate dehydrogenase (Doello et al., 2025). In contrast, the role of phosphorus reserves appears to vary among species. Compared to *Microcystis*, which has a high capacity for phosphorus uptake and storage (e.g., via polyphosphate bodies), phosphorus-sensitive genera such as *Dolichospermum* may experience limited by phosphorus scarcity (Wan et al., 2019).

### 4.3. A growing doubt: Is hormesis really beneficial?

Previous studies have often assumed that hormesis is beneficial for organisms adapting to their environment (Forbes, 2000; Parsons, 2001; Mushak, 2013). Our study further demonstrates the costs of hormesis (Cedergreen, 2008; Jager et al., 2013). This contradiction suggests that the beneficial nature of hormesis is not a universal rule but a context-dependent outcome that ought to be interpreted against the organism’s baseline physiological state. Our results demonstrated that a state of slow, normal growth can support a positive hormesis effect (**Fig. 2**). Conversely, hormesis is either irrelevant or mechanistically unjustified under two regimes: (i) near-optimal resource supply coupled with negligible stress; (ii) extreme resource scarcity (Liu et al., 2015a; Liu et al., 2015b). Species variation is another key factor; for instance, unicellular organisms may exhibit different hormetic responses compared to multicellular individuals (Feng et al., 2023). The latter may maintain a positive energy balance (Erofeeva, 2024) and demonstrate a more sustained positive response (Mushak, 2016). Moreover, in complex microbial systems where phytoplankton and bacteria interact closely (such as the phycosphere), mutual support is common (Kuhlisch et al., 2023). Bacteria can supply phytoplankton with essential nutrients, while phytoplankton reciprocate with organic carbon (You et al., 2025). This symbiotic exchange may help sustain a prolonged hormetic state (Lemos, 2026). Other possible cases are undoubtedly widespread in nature, and the tip of the iceberg of uncertainties regarding hormesis revealed by our study indicates that the threat of low-level chemical pollution demands careful evaluation.

### 4.4. Current limitations and implications for future environmental research

Given the vast diversity of chemicals and organisms, real-world risk assessment remains a major challenge. Organisms exhibit considerable trait variation (Schavemaker and Lynch, 2025), and the combined toxicological effects of different chemicals are virtually impossible to fully characterize (Yan et al., 2025). Our study focused on *Microcystis* and antibiotics as a representative species-pollutant system, and we believe that expanding this line of research will be valuable for addressing broader ERA challenges. The composition of organic constituents, including most emerging contaminants, in natural water bodies is highly complex and variable (Tanentzap and Fonvielle, 2024). The mechanisms underlying the actions of various organic components require substantially more research. In subsequent studies, we plan to identify organic components that cause shifts in *Microcystis* populations, including synthetic chemicals and naturally occurring or biologically secreted substances.

Notably, baseline growth rate is linked to hormesis via the dynamics of growth-related reserves (**Fig. 2**). The phenomenon also appears in antibiotic killing assays, where bacterial survival is governed by resource allocation that correlates with baseline growth rate (Bren et al., 2023). When growth is mildly limited, organisms store extra resources as reserves (Xiao et al., 2024). These easily accessible reserves then provide the energy that powers the overcompensation response (Gurney and Nisbet, 2004). This relationship suggests the applicability of the Dynamic Energy Budget (DEB) framework (Lemos, 2026), which provides a mechanistic basis for it (Feng et al., 2026). DEB framework is universally applicable across species and delineates the processing of various materials linking molecular-level metabolism to physiological functions at the whole-organism level (Marschmann et al., 2024). The DEB framework describes how organisms manage resources by absorbing, storing, and mobilizing matter and energy in reserves (Oliveira et al., 2026). The observed hormesis hinges on cells enhancing the mobilization of stored reserves to fuel the costs of physiological adjustments (Erofeeva, 2022), with a particular emphasis on shifting the balance of reserve allocation to direct more resources toward growth (Fig. 4a). This led to reduced investment in cellular structures and a depletion of the reserve pool. *Microcystis* exhibited DEB traits whereby it accumulated mobilizable unallocated reserves under slow-growth conditions to support hormetic responses, whereas at maximum growth rates these pools were almost depleted, limiting its ability to cope with mild stress (Fig. 4b) (Lemos, 2026).

**Fig. 4.**
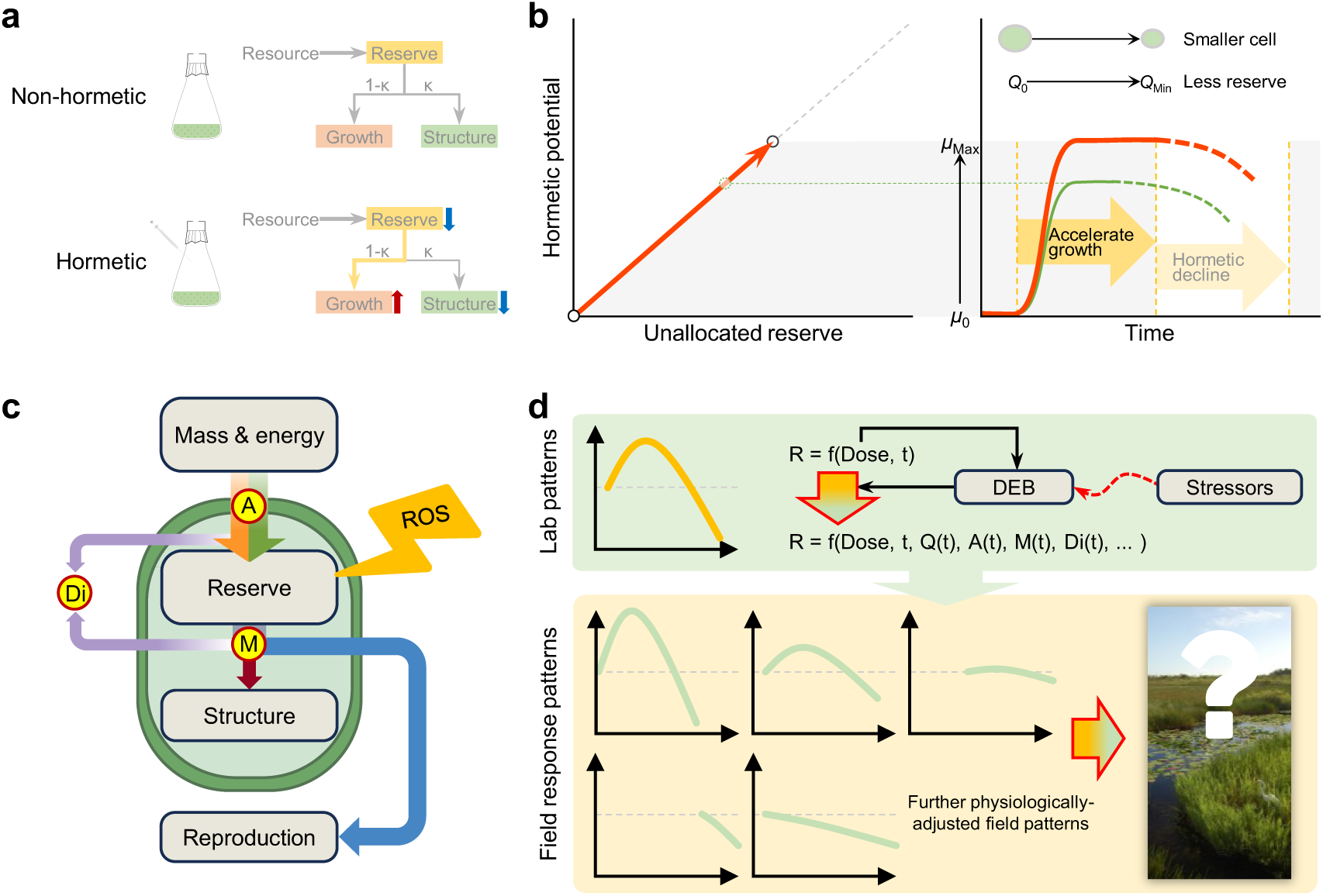
a. Schematic dynamic energy budget of cellular reserve flows. **b** Schematic relationship between unallocated reserves and potential promotion in growth rate. We speculate that hormesis may entail costs related to reserves. Hormesis can occur when cells utilize surplus reserves to support accelerated growth, albeit at the cost of depleting those reserves and decreasing cell size. When reserves are no longer sufficient to sustain the heightened growth, it will lead to a decline. Under this assumption, larger cellular reserves can support a stronger and more sustained hormesis effect (refer to the red and green curves in the panel, where their solid and dashed lines represent the rise and fall of the growth rate). **c** Schematic of Dynamic Energy Budget (DEB). **d** A forward-looking framework applies DEB theory to refine dose-response curves derived from controlled experiments, enabling extrapolation to real-world ecological contexts. *Q*_0_ and *Q*_Min_ represent the initial cellular quotas before stimuli and the minimum quotas at the end of exposure, respectively. *Q*_0_ - *Q*_Min_ denotes the unallocated reserve before exposure. *Q*_Min_ typically denotes the quota per cell to support the basical cell structure. *μ*_0_ represents the baseline growth rate before exposure. *μ*_Max_ represents the peak growth rate after exposure. Additional abbreviations: Assimilation (A); Dissipation (Di); Mobilization (M); Reactive oxygen species (ROS); Response (R); Exposure time (t). Panels **c** and **d** are adapted from our preprint (Feng et al., 2026). The wetland illustration in panel **d** was from Alexandra Fries, Integration and Application Network (https://ian.umces.edu/media-library), which is licensed under Attribution-ShareAlike 4.0 International (CC BY-SA 4.0) (https://creativecommons.org/licenses/by-sa/4.0/).

Nowadays, research on hormesis is approaching a critical stage for understanding its real-world toxicological implications (Veresoglou and Begum, 2024). Some ecosystem-related responses have been documented. For instance, changes in soil enzyme and microbial activities under metal stress in laboratory-treated samples (Zhang et al., 2023). These findings underscore the importance of translating extensive laboratory data, which constitute a valuable scientific resource, into reliable real-world predictions (Sun and Wu, 2026). Pollutant effects in organisms where assimilation clearly exceeds dissipation may be predicted using dose-response curves similar to those established under ideal laboratory conditions. However, such conditions are seldom observed in single-celled microorganisms (Mushak, 2016). Hence, microbial hormesis in nature likely represents the most formidable obstacle to advancing the risk prediction of hormesis. Predictions under environmental stress can be refined by adjusting the maximum stimulatory response and the width of the hormetic zone (see Agathokleous, 2025). This can be achieved by tuning DEB model parameters to modify the dose-response curve function (Fig. 4d), which includes establishing boundary conditions under resource limitations. For example, under strict nitrogen-limited conditions, the promotional effect on growth is nearly eliminated (**Fig. 2**). A hormetic effect is also unlikely to occur when the trait cost or total energy budget of an organism exceeds its environmental fitness (Schavemaker and Lynch, 2025; Lemos, 2026). Adjusting the parameters of dose-response curves based on physiological status, especially when exploring thresholds, may uncover broader research frontiers in toxicology. Furthermore, in ecological studies, particularly those involving population-level dynamic simulations, it is important to advance the understanding of hormesis and other toxicological mechanisms within model organism-contaminant systems and extend these insights to the aquatic ecosystem scale. Overall, this study highlights the feasibility of extrapolating laboratory-derived dose-response patterns, corrected for key physiological traits, to predict pollutant impacts in aquatic ecosystems. It further emphasizes that accounting for cellular reserves and resource allocation strategies in ecotoxicity assessments will enable a more accurate reflection of real-world effects of emerging contaminants.

## 5. Conclusion

This study shows that baseline growth rates and the dynamics of cellular reserves are significantly associated with the potential for hormesis in harmful cyanobacterium *Microcystis*. Slow-growing populations experience a substantial increase in growth rate under low-intensity antibiotic stress, albeit at the cost of disrupting stoichiometric homeostasis, as evidenced by a rapid reduction in cellular carbon and nitrogen quotas. In contrast, fast-growing groups exhibit limited sensitivity to stress, likely due to their limited unallocated reserves. Thus, a key source of uncertainty in hormesis effects is the physiological state, especially the mobilization of reserves. In real-world aquatic environments, our study highlights that accurately predicting the impacts of hormesis requires accounting for both physiological states and environmental conditions. Based on these findings, we identify two research priorities: (i) developing toxicity threshold models based on energy budgets and refining toxicity prediction methods for real-world scenarios and (ii) advancing mechanistic studies in model organism-contaminant pairs and extending them to natural water environments.

## Acknowledgments

The authors wish to express their deepest gratitude to Dr. Jolanda M.H. Verspagen of the University of Amsterdam for generously dedicating her time and offering insightful comments that significantly improved the clarity of this work. The authors acknowledge DeepSeek (https://www.deepseek.com/) for its assistance in language polishing. Furthermore, we thank the ZhuWei Research Team at Hohai University for their kind provision of the field water samples. This work was funded by the National Natural Science Foundation of China (No. 32301373) and the Research Startup Fund of Jiaxing University.

## CRediT authorship contribution statement

**Ganyu Feng:** Conceptualization, Methodology, Investigation, Data Curation, Writing - Original Draft, Writing - Review & Editing, Visualization, and Funding acquisition. **Zhipeng Duan:** Methodology, Validation, Formal analysis, Funding acquisition, and Writing - Review & Editing. **Changwei Hu:** Writing - Review & Editing. **Juan Wang**: Writing - Review & Editing. **Jie Xu**: Writing - Review & Editing. **Yuejin Zhang:** Writing - Review & Editing, Resources, Supervision. **Xiang-Zhou Meng:** Methodology, Supervision, Writing - Review & Editing, and Project administration.

## Data availability

Data are available from the authors upon reasonable request.

