## Supplemental Files for "Baseline growth rate shapes pollutant impacts in aquatic environments: Evidence from antibiotic responses in the harmful cyanobacterium *Microcystis*"

**Appendix S1. Supplementary Texts................................................................Page S3**

**Text S1.** Preparation of the stock solution of solid phase extraction.

**Text S2.** Calculation of specific growth rate, absolute response, and relative response for the entire duration of exposure.

**Text S3.** Transcriptomic analysis.

**Appendix S2. Supplementary Figures...........................................................Page S7**

**Fig. S1.** Relative response and absolute response under antibiotics in *Microcystis* sp. analyzed using data collected from a shared dataset. Relations between the baseline growth rate during steady stable growth stage and the dilution rates of continues culture and semi-continuous culture. Variation of the cell concentration and growth rate with exposure time.

**Fig. S2.** Changes in cell concentration before and during exposures. Absolute response values at specific time points.

**Fig. S3.** Cell states before, after, and at the end of exposures.

**Fig. S4.** Changes in cellular dry weight and phosphorus content at various dilution rates.

**Fig. S5.** Exposures to antibiotics in modified BG11 media.

**Fig. S6.** The growth process of *Microcystis* exposed to 200 ng $L^{-1}$ antibiotic under dark conditions.

**Fig. S7.** Heatmap of expression levels of all mapped genes.

**Fig. S8.** Volcano plots of statistical significance against $\log_2$(FC) of DEGs.

**Fig. S9.** Scatter plots showing $\log_2$(FC) values for shared DEGs.

**Fig. S10.** Gene expression of DEGs enriched in the KEGG pathways.

**Fig. S11.** Heatmaps for photo-related genes.

**Fig. S12.** Schematic diagram of the experimental process and specific growth rate.

**Appendix S3. Supplementary Tables...........................................................Page S19**

**Table S1.** Details of linear fitting results.

**Table S2.** Residual levels of dissolved inorganic carbon, nitrogen, and phosphorus in the culture media at the end of the exposure periods.

**Table S3.** Expression levels in RPKM of the 105 DEGs.

**Table S4.** Enriched KEGG pathways.

**Supplementary References..................................................................Page S29**

### Appendix S1. Supplementary Texts

#### *Text S1. Preparation of the stock solution of solid phase extraction (SPE)*

A 500 mL water sample was collected on December 25, 2024, from the water surface of Lake Taihu, China (at 120.18552670 °E, 31.35260487 °N). The sample was then transported to the laboratory via S.F. Express in a cool, dark environment using ice as soon as possible. The water sample was filtered through a clean 0.45-μm glass fiber filter, and the filtrate was extracted using an SPE system. Initially, the pH was adjusted to 3.0 with 2 mol $L^{-1}$ HCl. Samples were loaded at a flow rate of 3 – 5 mL $min^{-1}$ onto Waters Oasis HLB cartridges (500 mg, 6 mL), which had been preconditioned with 5 mL of methanol and 5 mL of ultrapure water (pH 3.0). The cartridges were then rinsed with 5 mL of a 5% methanol aqueous solution and 5 mL of a 1% methanol solution, and dried under vacuum for 15 min. Organic contaminants were eluted with methanol. The eluent was then concentrated under a gentle nitrogen stream to near dryness, after which it was dissolved in 2 mL of methanol and stored at -20°C as a stock solution. The day before exposure, 0.5 mL of the stock solution was concentrated under a gentle nitrogen stream to near dryness, and then redissolved in 2.5 mL of ultrapure water. This solution was the SPE stock for exposure experiment and stored overnight at 4°C. Consequently, the enrichment of the SPE stock was 50 times, and 50 μL of the SPE stock was added to each test tube of the treatments. The axenic stock solution was prepared before use by passing it through sterile filters with a pore size of 0.22 μm.

#### *Text S2. Calculation of specific growth rate ($\mu_{Ent.}$), absolute response (AR), and relative response (RR) for the entire duration of exposure*

The daily specific growth rates ($\mu_i$) was averaged (denoted as $\mu_{Ent.}$). A fitted line was selected only if a linear regression applied to the natural logarithm-transformed cell concentrations from the start to the end of exposure yielded at least three data points. Among all qualifying lines, the one with the highest $R^2$ value was chosen. If no data set had an $R^2$ value greater than 0.9, we fitted the data from 0 hours to 72 hours. In cases where there was a decrease in cell concentration, we fitted the data from the end of monitoring to the start of exposure to determine the data points participating in the calculation of $\mu_{Ent.}$, following the same principles as described above. The absolute response (AR) and relative response (RR) for the entire duration of exposure were calculated as follows:

$$\mu_i = \ln\left(\frac{N_i}{N_{i-24}(1-DR)}\right) \quad (S1)$$

$$\mu_{Ent.} = \frac{24}{k}\sum_0^k \mu_i \quad \text{if cell concentration increases} \quad (S2)$$

$$\mu_{Ent.} = \frac{24}{m-k}\sum_k^m \mu_i \quad \text{if cell concentration decreases} \quad (S3)$$

$$RR = (\mu_{Ent.} - \mu_0)/\mu_0 \times 100\% \quad (S4)$$

$$AR = \mu_{Ent.} - \mu_0 \quad (S5)$$

where, $\mu_i$ and $N_i$ are the specific growth rate and cell concentration after 'i' hours of exposure onset, respectively. During the exposure experiment, cell concentration was measured every 24 hours. $\mu_0$ is the baseline growth rate before exposure, which depends on the dilution rate (refer to **Figure S1c**). The time spans '0 – k' and 'k – m' correspond to the data sets resulting in the highest $R^2$ value when cell concentration increases and decreases, respectively.

***Text S3. Transcriptomic analysis***

Transcriptomes from two exposure experiments, conducted with a dose of 200 ng $L^{-1}$ and different dilution rates of 0.05 and 0.20 $day^{-1}$, were sampled and then analyzed by a sequencing company (Biozeron Biotech Co., Ltd., Shanghai, China):

Cells from 4 mL samples (from cultures at a dilution rate of 0.05 $day^{-1}$) and 15 mL samples (from cultures at a dilution rate of 0.20 $day^{-1}$) were collected, both the day before and 48 hours after exposure, as well as at the end of exposure. The cells were pelleted by centrifugation at 10000 *g* for 15 min (4°C), and then the samples were sent to the sequencing company on dry ice.

Total RNA was extracted from the samples using TRIzol® Reagent (Invitrogen, CA, USA) following the manufacturer's instructions. Genomic DNA was removed using DNase I (TaKaRa, Shiga, Japan). The purity of the RNA and the integrity of the samples were assessed using a Nanodrop2000 spectrophotometer (NanoDrop Technologies, MA, USA) and an Agilent Bioanalyzer 2100 system (Agilent Technologies, CA, USA), respectively. High-quality RNA sample (OD260/280=1.8~2.2, OD260/230≥2.0, RIN≥6.5, 28S:18S≥1.0, >10 μg) is used to construct sequencing library.

RNA-seq strand-specific libraries were prepared following the TruSeq RNA sample preparation Kit from Illumina (San Diego, CA, USA), using 5 μg of total RNA. Briefly, rRNA removal was performed using the RiboZero rRNA removal kit (Epicenter, WI, USA), and the RNA was then fragmented using fragmentation buffer. Subsequently, cDNA synthesis, end repair, A-base addition, and ligation of the Illumina-indexed adaptors were carried out according to Illumina's protocol. The libraries were size selected for cDNA target fragments of 200 – 300 bp on 2% Low Range Ultra Agarose, followed by PCR amplification using Phusion DNA polymerase for 15 PCR cycles. After quantification using TBS380, the paired-end libraries were sequenced using the Illumina NovaSeq 6000 sequencing platform (150bp*2, Shanghai BIOZERON Co., Ltd). The sequencing dataset has been uploaded to the Sequence Read Archive (SRA) database at the National Center for Biotechnology Information (NO. PRJNA1153622; https://www.ncbi.nlm.nih.gov/sra/?term=PRJNA1153622).

The raw paired-end reads were trimmed and quality controlled using Trimmomatic with the parameters (SLIDINGWINDOW:4:15 MINLEN:75) (version 0.36). The clean reads were then aligned to the reference genome with the orientation mode using Rockhopper. Rockhopper is a comprehensive and user-friendly system specifically designed for computational analysis of bacterial RNA-seq data. It takes RNA sequencing reads generated by high-throughput sequencing technology as input. The software was used to calculate gene expression levels with default parameters.

The clean rates of all samples in this study ranged from 96.30% to 97.25%, and the rRNA rates were all less than 2.94%. *Microcystis aeruginosa* PCC 7806 (https://www.ncbi.nlm.nih.gov/datasets/genome/GCF_030553035.1/) was selected as the reference genome for further data analysis, and the mapped ratios of all samples ranged from 81.53% to 94.81%.

To identify differentially expressed genes (DEGs) between two samples, the expression level for each transcript was calculated using the reads per kilobase per million mapped reads (RPKM) method. The sequencing company performed the DEGs analysis using edgeR. The DEGs were selected based on the following criteria: genes with a Benjamini-Hochberg adjusted *p* value ($P_{adj}$) $\leq 0.05$ and $|\log_2 FC| \geq 1$. Here, FC represents fold change. KEGG pathway analysis was conducted using KOBAS 3.0.

### Appendix S2. Supplementary Figures

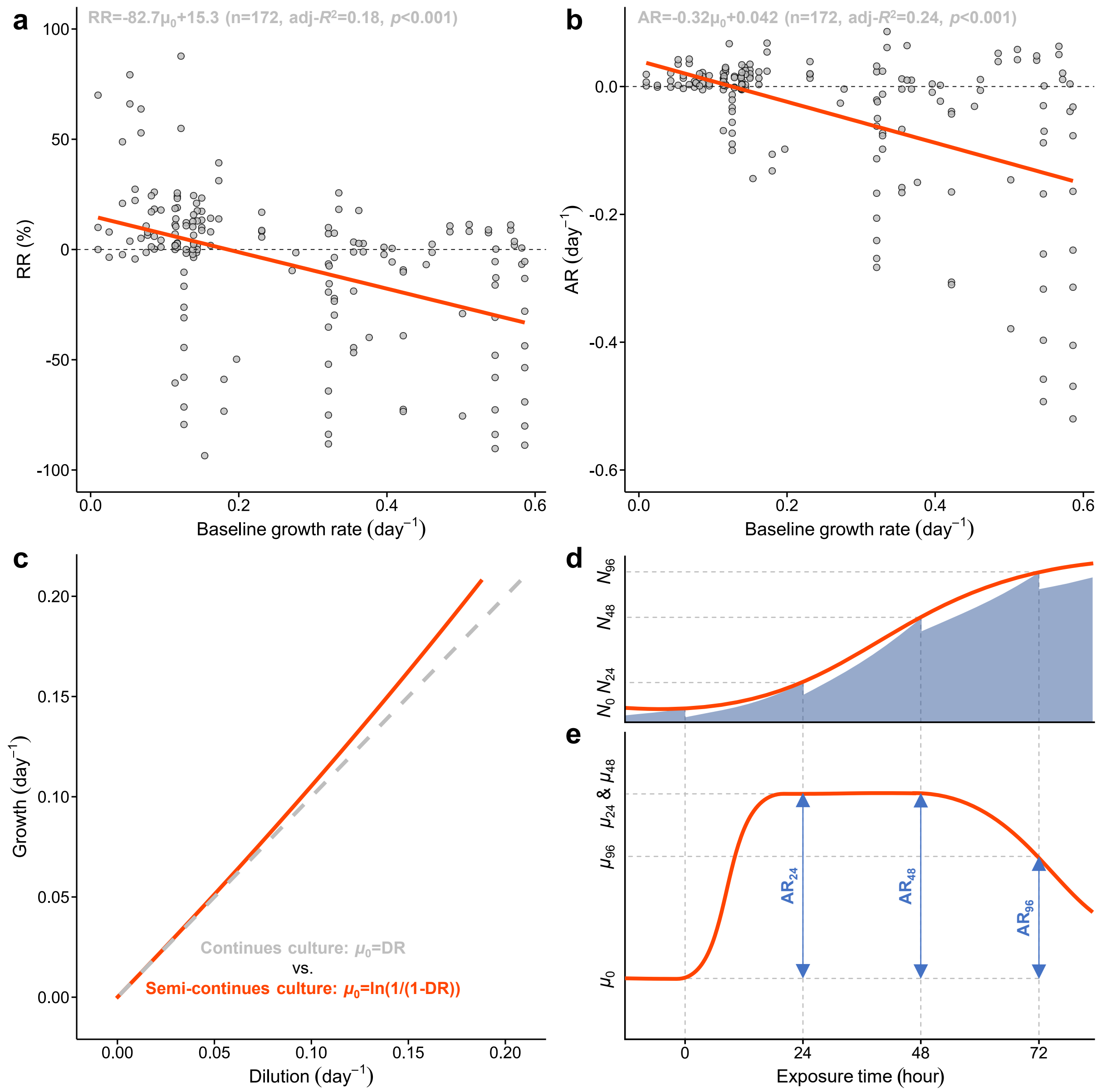


**Fig. S1.** Relative response (**a**: RR) and absolute response (**b**: AR) of specific growth rate under antibiotics in microalgae analyzed using data collected from a shared dataset (https://cstr.cn/31253.11.sciencedb.08733; "Data04 (Antibiotic_Colony_vs_sjngle cell).xlsx" of Version 3) of a previous publication (Feng et al., 2023). Data with RRs ranging from -100% to 100% and antibiotic concentrations ranging from 1 ng $L^{-1}$ to 1 mg $L^{-1}$ were plotted. The mean and median antibiotic concentrations were 22.6 μg $L^{-1}$ and 200 ng $L^{-1}$, respectively. The AR at a specific time point was calculated by subtracting the baseline growth rate ($\mu_0$) from the growth rate at that moment (Eq. 2). RR is defined as the ratio of the change in growth rate to the baseline growth rate, expressed as a percentage (Eq. 3). The red lines in panel **a** and **b** were fitted by the "ggplot2" package of R. Results show that microalgae with lower baseline specific growth rates have a higher percentage of positive responses (i.e., RR > 0 or AR > 0) than other groups with higher baseline growth rates. **c** illustrates the relations between the baseline growth rate during steady stable growth stage and the dilution rates (DR) of continues culture and semi-continuous culture (with a daily dilution frequency). The baseline growth rate of continuous culture is the same as the dilution rate during the steady state growth stage. For semi-continuous culture, the growth rate from the time of dilution on one day (i.e., after dilution) until the next dilution is calculated based on the increase in cell concentration. During this period, growth compensates for the loss in cell concentration caused by dilution, resulting in a stable phenomenon where the observed cell concentration remains consistent each time. Therefore, the growth rate of semi-continuous culture is also related to the dilution rate. Under our once-daily dilution frequency, the growth rate is the natural logarithm of the value obtained by dividing the baseline biomass by the biomass after dilution. **d** shows variation of the cell concentration ($N$) with exposure time (0, 24, 48, 96, ... hours). $N_0$ represents the baseline cell concentration of semi-continuous culture. Semi-continuous cultures can be viewed as a series of short-cycle batch experiments, where each new batch starts right after the previous one ends (LaRoche et al., 2010). Therefore, the actual change in cell concentration corresponds to the shaded area, while the red curve depicts the smooth trend in cell concentration change based on the measured cell concentration. **e** shows variation of the growth rate with exposure time. The curves are for illustrative purposes only and does not indicate an actual trend. Before exposure, the cellular growth rate was at the baseline level.

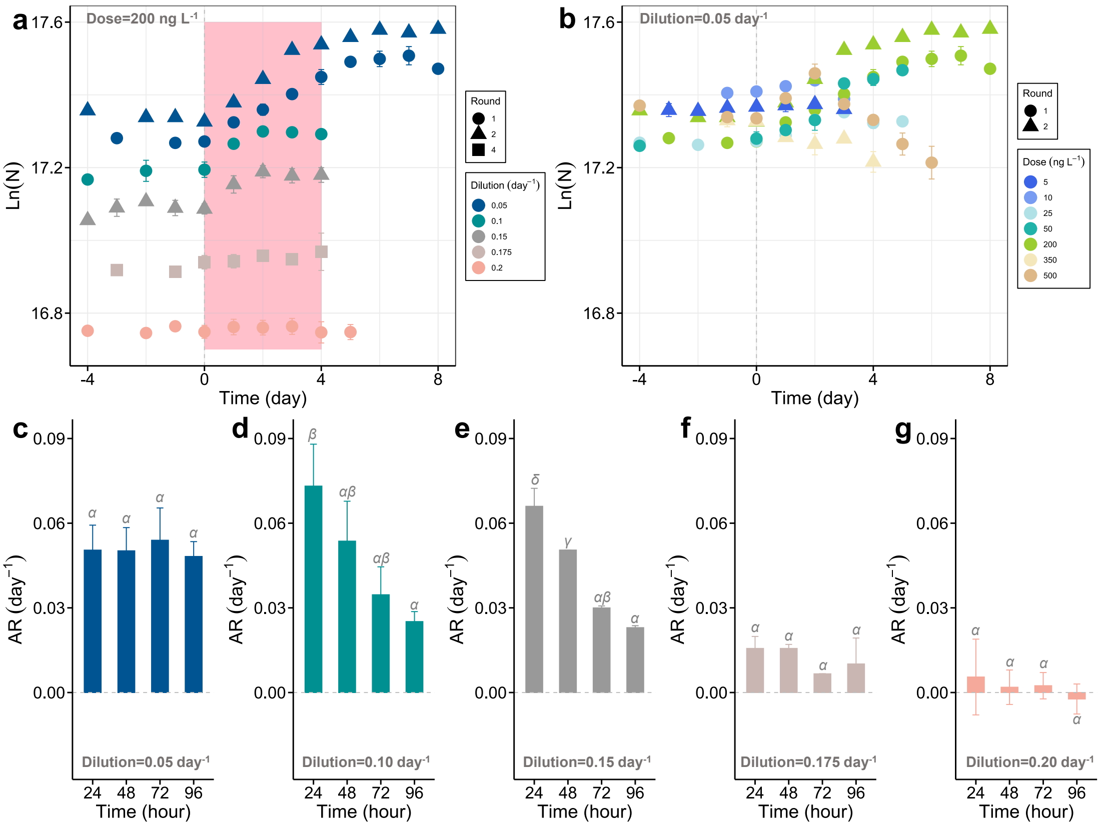


**Fig. S2.** Cell concentration (N; cells mL$^{-1}$) before and during exposures presented for different dilution rates (**a**) and antibiotic doses (**b**). Each exposure incubation was prepared with two biological replicates. Panels **c** to **g** display the absolute response (AR) values at specific time points, which are derived from the data in the pink shaded area of panel **a** and calculated using Eq. (1 – 2). Greek letters (α, β, and γ) indicate a significant difference ($p < 0.05$) between time points. In Round 1, we conducted exposure experiments at dilution rates of 0.05 day$^{-1}$ (dose = 10, 25, 50, 200, and 500 ng L$^{-1}$), 0.10 day$^{-1}$ (dose = 200 ng L$^{-1}$), and 0.20 day$^{-1}$ (dose = 200 ng L$^{-1}$). In Round 2, we tested three doses (5, 200, and 350 ng L$^{-1}$) at a dilution rate of 0.05 day$^{-1}$, with the 200 ng L$^{-1}$ exposure serving as a repeated experiment, and additionally performed exposures at 200 ng L$^{-1}$ with a dilution rate of 0.15 day$^{-1}$. Round 4 included experiments at 200 ng L$^{-1}$ and 0.175 day$^{-1}$.

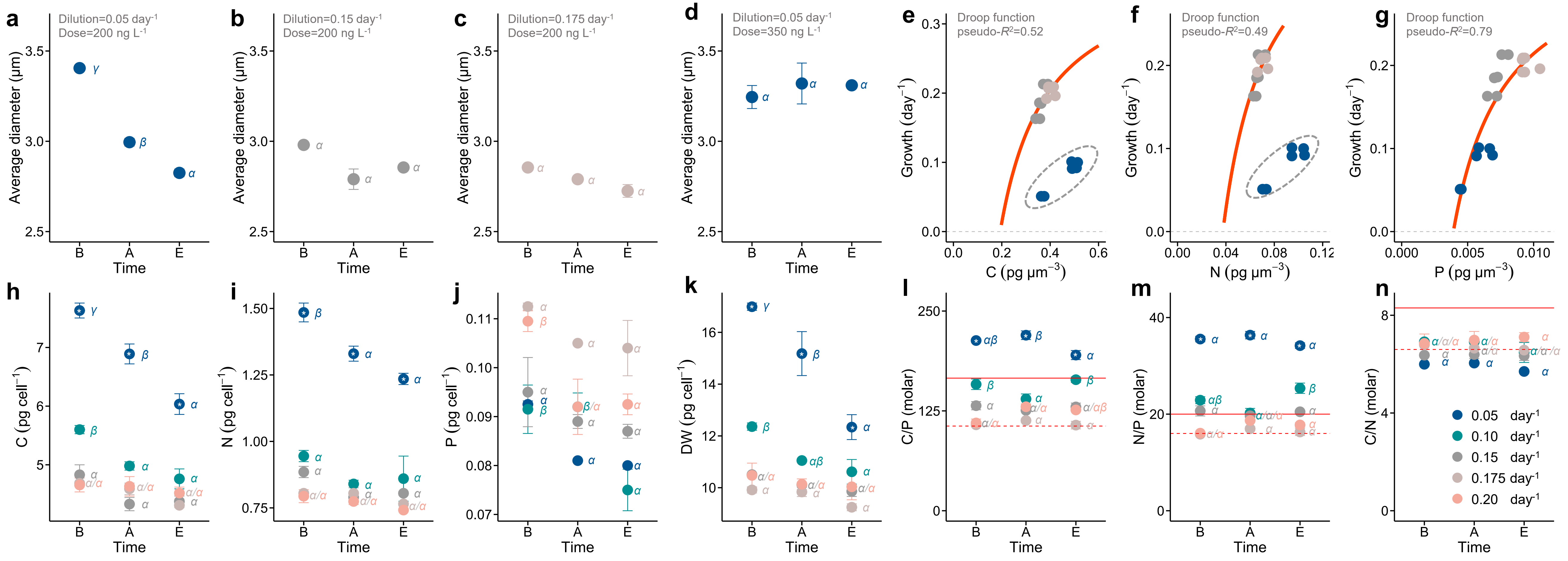


**Fig. S3.** Cell states before ('B'), 48 h after ('A'), and at the end ('E') of exposures: cell diameter (**a-d**), cellular carbon (**h**), nitrogen (**i**) and phosphorus (**j**) contents, dry weight (**k**), and elemental molar ratios (**l-n**; dashed lines indicate Redfield ratio (Klausmeier et al., 2004; Seelen et al., 2025) and solid lines indicate Sterner ratio(Andersen et al., 2025)). Panels **e-g** show curve fittings between the specific growth rate and element content per unit cell volume according to the Droop equation (Droop, 1973). Here, the diameters of at least 200 cells per biological sample were measured at a magnification of 400 × (Olympus microscope, Tokyo, Japan), and the cell volume was calculated using the formula for the volume of a sphere. The data points circled by dashed lines were not included in the fitting. Greek letters (α, β, and γ) indicate a significant difference ($p < 0.05$) between time points. The asterisk (*) indicates significant differences at the $p < 0.05$ level between the lowest dilution rate and all other dilution rates. All data were derived from exposure experiments based on standard BG11 medium.

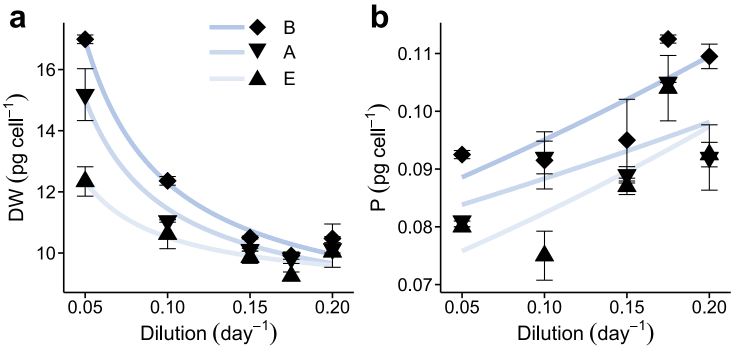


**c**

| | Time node | Function | a | b | pseudo-$R^2$ |
|---|---|---|---|---|---|
| | B | | 0.47 | 7.6 | 0.98 |
| DW | A | $Q$=a/DR+$Q_{Min,\infty}$ | 0.36 | 7.9 | 0.96 |
| | E | | 0.18 | 8.7 | 0.86 |
| | B | | 0.20 | 3.5 | 0.99 |
| C | A | $Q$=a/DR+$Q_{Min,\infty}$ | 0.16 | 3.5 | 0.94 |
| | E | | 0.11 | 3.7 | 0.94 |
| | B | | 0.046 | 0.55 | 0.98 |
| N | A | $Q$=a/DR+$Q_{Min,\infty}$ | 0.037 | 0.55 | 0.95 |
| | E | | 0.033 | 0.57 | 0.97 |
| | B | | 0.083 | 1.4 | 0.63 |
| P | A | $Q$=a*e$^{(DR*b)}$ | 0.080 | 1.1 | 0.42 |
| | E | | 0.070 | 1.7 | 0.55 |

**Fig. S4.** Changes in cellular dry weight (**a**; DW) and phosphorus content (**b**; P) at various dilution rates. 'B', 'A', and 'E' denote samples taken before exposure, 48 hours after exposure, and at the end of the experiment, respectively. **c** is the details of curve fitting results. '$Q$' is one of indicators (i.e., cellular quota of carbon, nitrogen, or phosphorus), $Q_{\mathrm{Min},\infty}$ is the minimum cellular quota of carbon or nitrogen as the growth rate approaches infinity, 'DR' is the dilution rate, and 'a' and 'b' are constants. The sample size in each fitting group was 10.

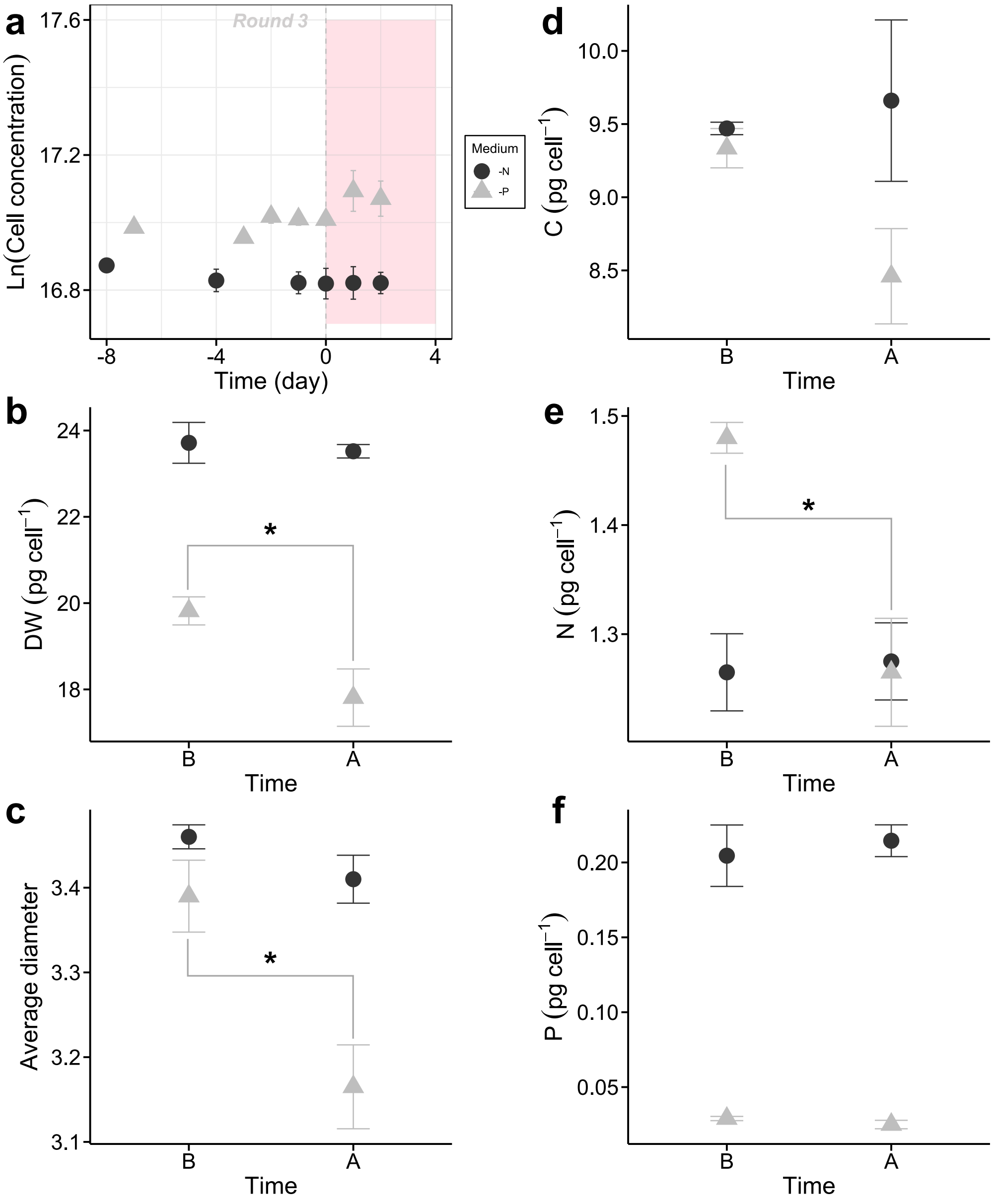


**Fig. S5.** Exposures to antibiotics in modified BG11 media (-N and -P). **a** Changes in cell concentration (cells $mL^{-1}$) before and during exposures. **b** Dry weight (DW) before ('B') and after ('A') exposures. **c** Average cell diameter. The diameters of at least 200 cells per biological sample were measured at a magnification of 400 × (Olympus microscope, Tokyo, Japan). **d-f** Cellular element contents of carbon (C), nitrogen (N), and phosphorus (P). The asterisk (*) indicates significant differences at the $p < 0.05$ level before and after exposure.

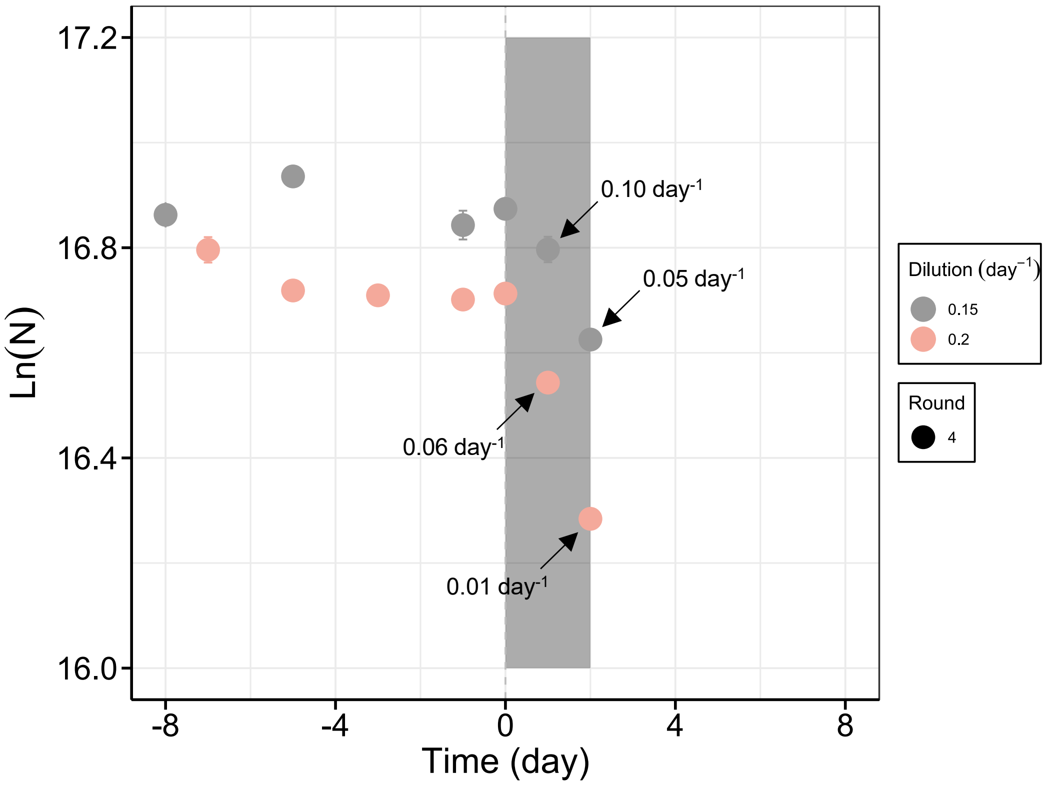


**Fig. S6.** The growth process of *Microcystis* exposed to 200 ng $L^{-1}$ antibiotic under dark conditions. The shaded area indicates the light-deprived phase. The results indicate that after blocking the light source, *Microcystis* did not exhibit an hormetic response (compared with **Fig. S2**a). The growth rate significantly decreased and continued to decline as the dark period extended, while biomass diminished due to dilution. Compared to the fast dilution group (0.20 $day^{-1}$), the net growth rates were comparatively higher under the slower dilution condition (0.15 $day^{-1}$). The above results highlight the impact of energy supply on the hormesis effect: the slowly growing group possesses more cellular reserves to buffer against the decrease in growth rate.

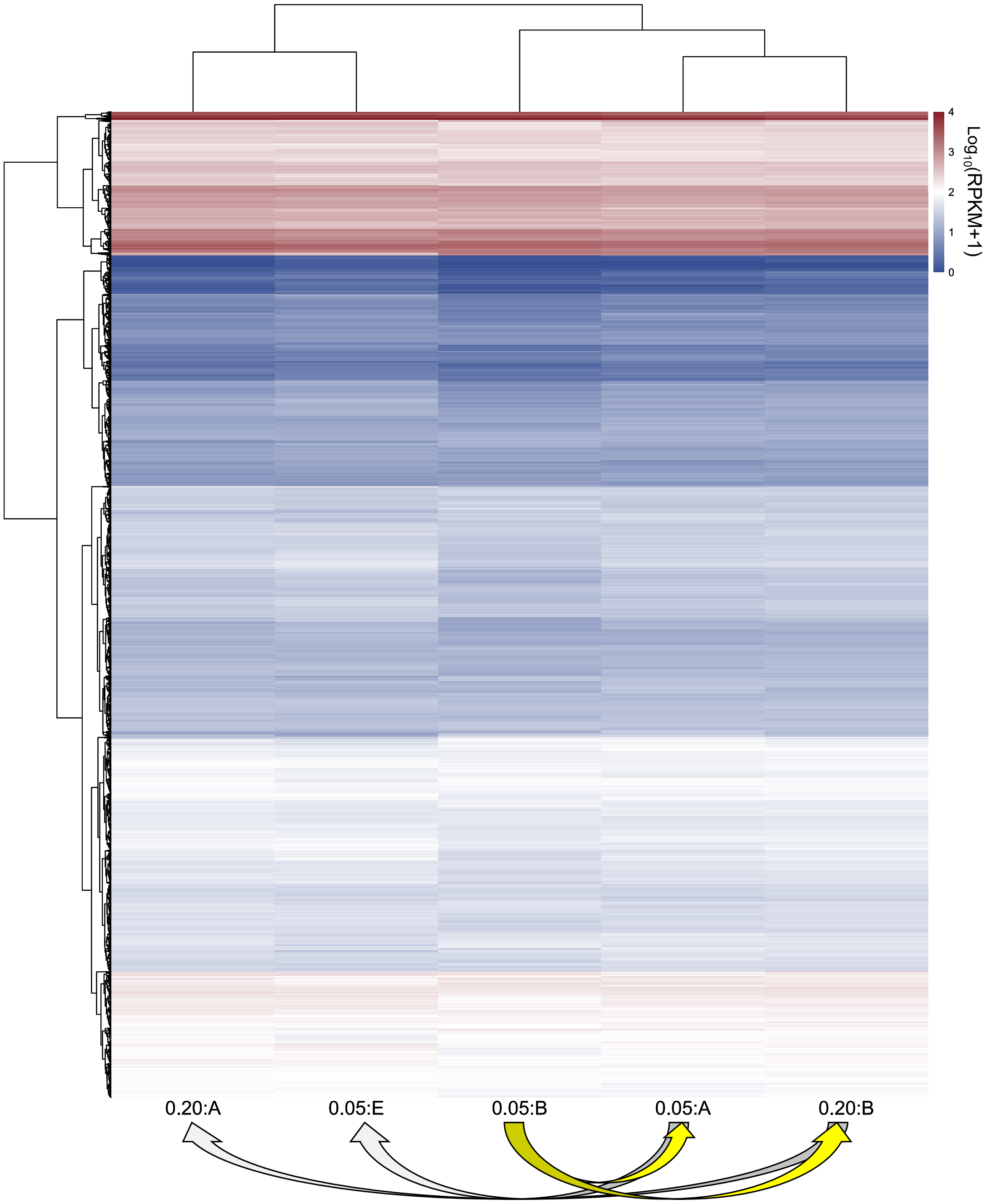


**Fig. S7.** Heatmap of expression levels in $\log_{10}$(RPKM+1) of all mapped genes.

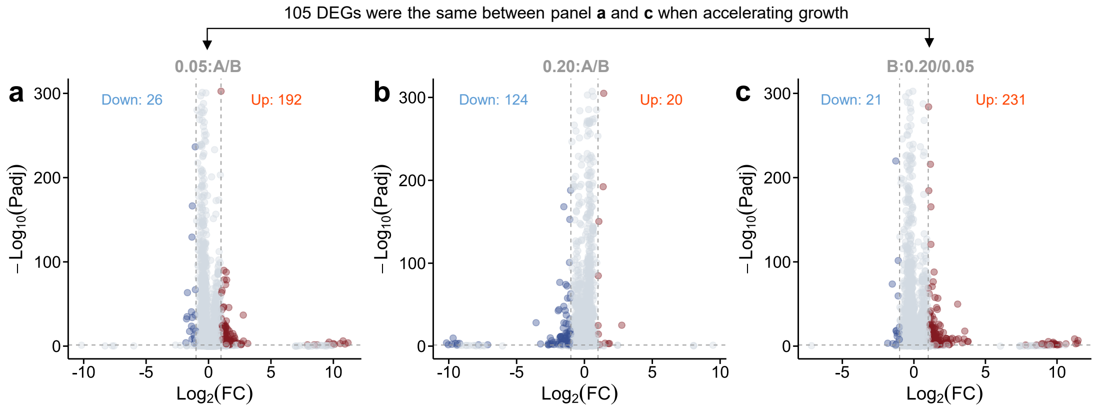


**Fig. S8.** Volcano plots of statistical significance ($-\log_{10}$Padj) against $\log_2$(FC) of DEGs (Padj $\leq$ 0.05) in 0.05:A/B, 0.20:A/B, and B:0.20/0.05 comparisons, respectively.

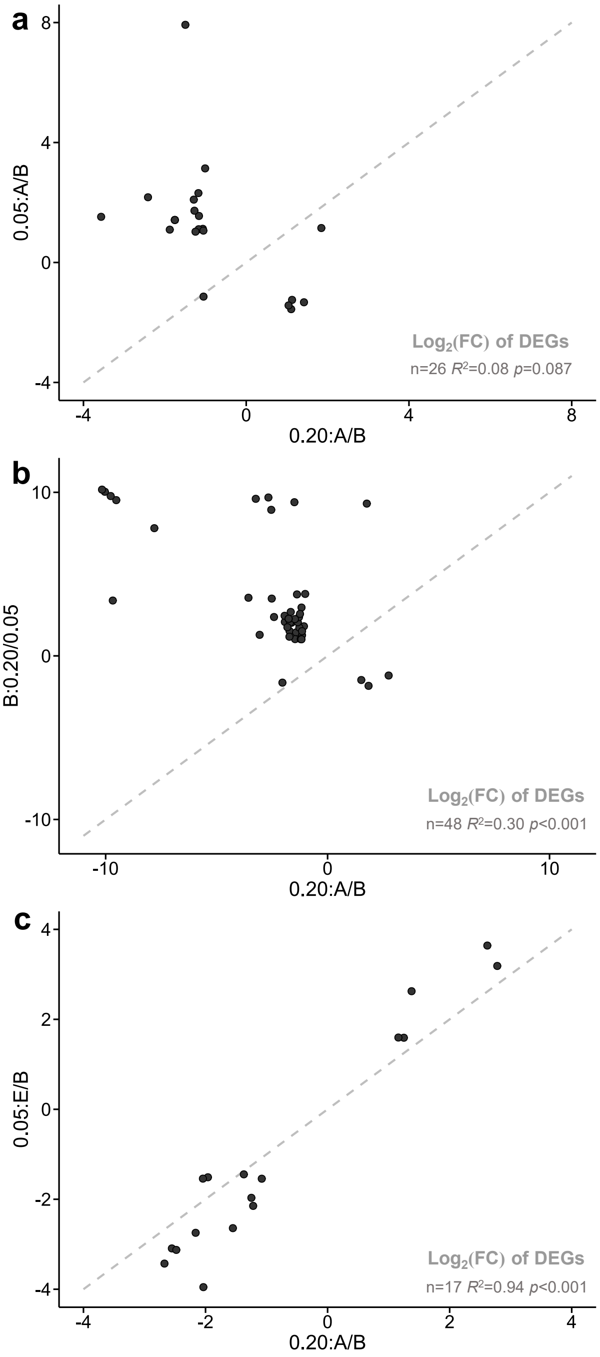


**Fig. S9.** Scatter plots showing $\log_2$(FC) values for shared DEGs under different conditions in **Fig. 3**d: (**a**) between 0.05:A/B and 0.20:A/B, (**b**) between B:0.20/0.05 and 0.20:A/B, and (**c**) between 0.05:E/B and 0.20:A/B. For each panel, the best-fit line's results are provided (n, $R^2$, and $p$; the fitted lines are not drawn on the panels). A dashed line with a slope of 1 and an intercept of 0 is included as a reference to indicate the expected 1:1 relationship.

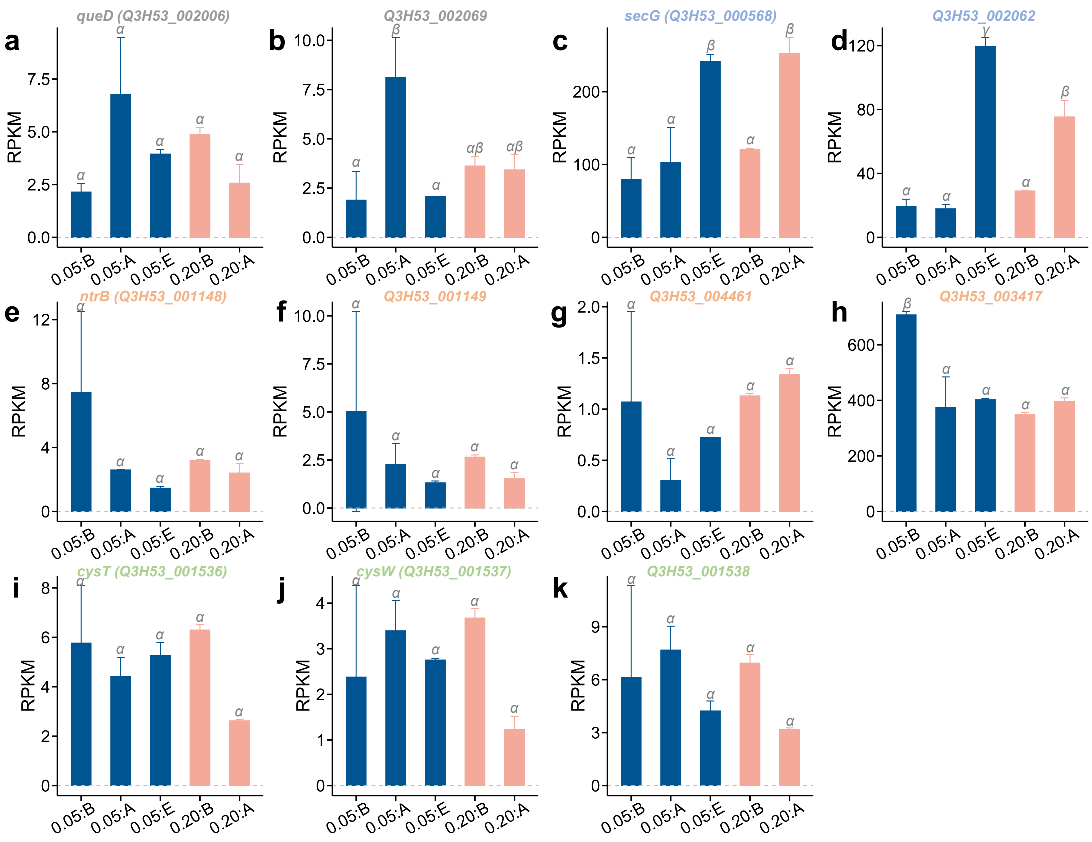


**Fig. S10.** Gene expression of DEGs enriched in the KEGG pathways listed in **Table S4**. Greek letters (α, β, and γ) indicate significant differences ($p < 0.05$).

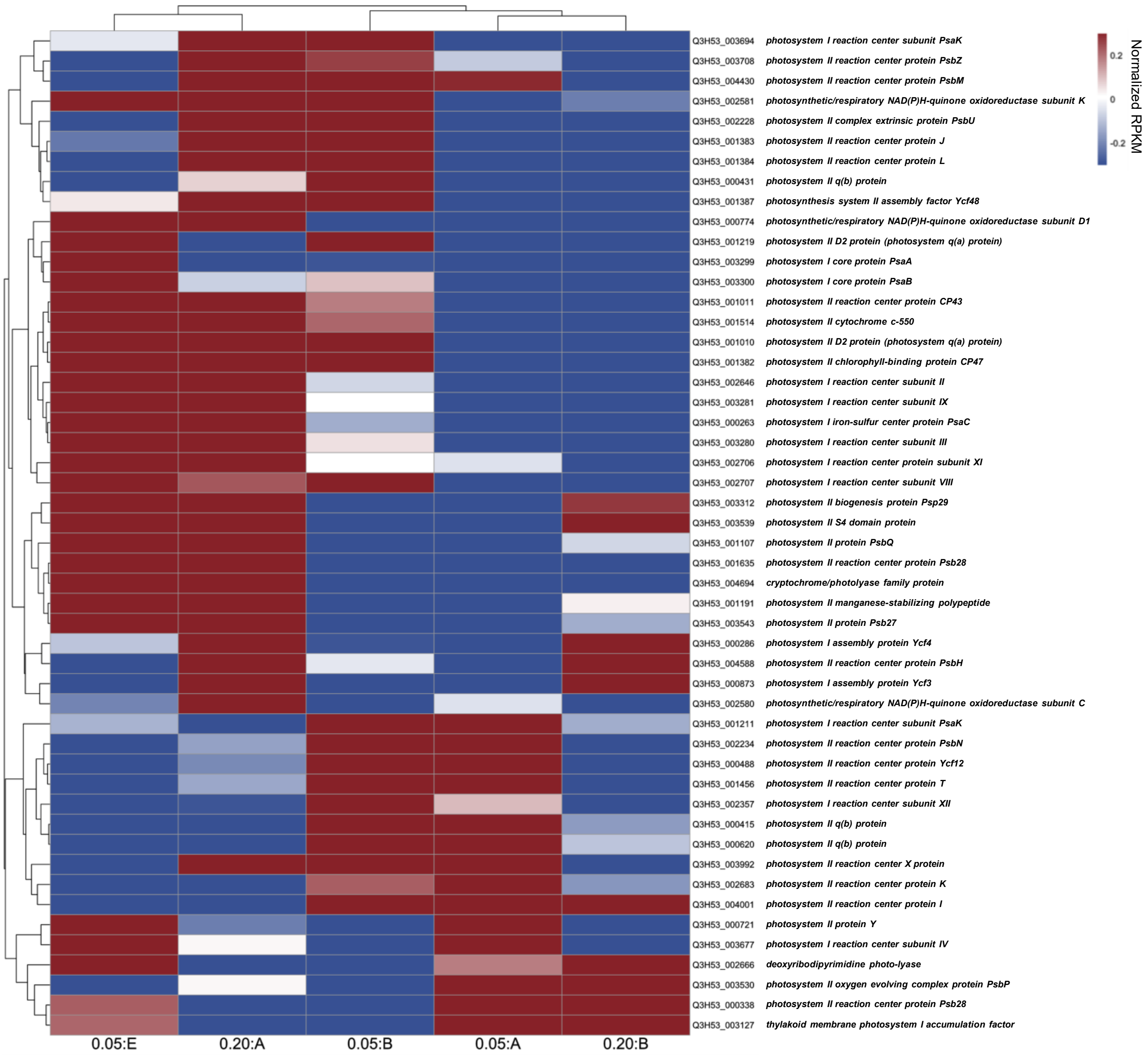


**Fig. S11.** Heatmaps for photo-related genes using the ‘pheatmap’ package with scale = ‘row’.

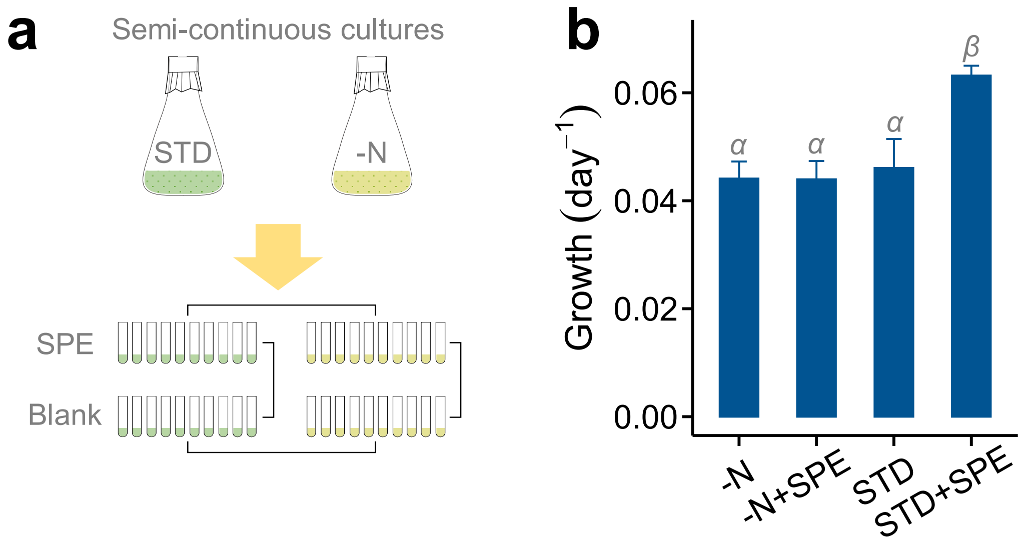


**Fig. S12.** **a** Schematic diagram of the experimental process for semi-continuous cultures and exposures. **b** Specific growth rate of four groups (-N, -N+SPE, STD, and STD+SPE). Greek letters (*α* and *β*) indicate a significant difference ($p < 0.05$).

**Appendix S3. Supplementary Tables**

**Table S1.** Details of linear fitting results. The fitting formula is y=a*x+b, where 'y' represents the absolute response after 48 hours of exposure onset ($AR_{48}$), 'x' is one of indicators, and 'a' and 'b' are constants. The sample size (n) in each fitting group was 10.

| **Indicators** | **a** | **b** | **adj-$R^2$** | ***p*** |
|---|---|---|---|---|
| $DW_{0\text{-}48}$ | 0.018 | 0.019 | 0.28 | 0.07 |
| $C_{0\text{-}48}$ | 0.063 | 0.0084 | 0.78 | 0.0003 |
| $N_{0\text{-}48}$ | 0.28 | 0.012 | 0.62 | 0.004 |
| $P_{0\text{-}48}$ | -1.8 | 0.049 | 0.25 | 0.08 |

**Table S2.** Residual levels of dissolved inorganic carbon (DIC), nitrogen (DIN), and phosphorus (DIP) in the culture media (STD, -N, and -P) at the end of the exposure periods.

| Dilution ($day^{-1}$) | Medium | DIC ($mg\ L^{-1}$) | DIN ($mg\ L^{-1}$) | DIP ($mg\ L^{-1}$) |
|---|---|---|---|---|
| 0.05 | STD | 7.72±1.05 | 174.13±13.77 | 3.012±0.241 |
| 0.20 | STD | 8.84±0.86 | 206.56±5.41 | 4.683±0.404 |
| 0.05 | -N | 7.68±0.88 | 0.29±0.04 | 3.084±0.075 |
| 0.05 | -P | 13.21±0.96 | 230.82±7.64 | 0.031±0.004 |

**Table S3.** Expression levels (mean ± standard deviation) in RPKM of the 105 DEGs common to groups 0.05:A/B and 0.20:B/0.05. The heatmap provides a comparison among groups at the level of each gene, which was drawn using Excel (Microsoft Office 2016). The warmer the color, the larger the value among groups.

| **GeneID** | **0.05:B** | **0.05:A** | **0.20:B** | **0.20:A** | **0.05:E** | **Product** |
|---|---|---|---|---|---|---|
| *Q3H53_000023* | 27.53±4.94 | 58.23±6.95 | 60.1±0.13 | 55.05±4.37 | 79.95±0.53 | type II toxin-antitoxin system RelE/ParE family toxin |
| *Q3H53_000059* | 5.93±1.14 | 12.97±0.24 | 12.73±0.21 | 7.5±0.77 | 10.61±0.35 | DUF5615 family PIN-like protein |
| ***Q3H53_000108*** | **335.88±24.65** | **147.33±45.81** | **160.47±4.62** | **239.37±2.21** | **259.53±6.25** | **nitrate ABC transporter permease** |
| *Q3H53_000201* | 6.74±2.32 | 15.72±2.89 | 21.52±0.95 | 23.1±7.17 | 46.46±1.52 | hypothetical protein |
| *Q3H53_000214* | 2.48±0.21 | 5.03±0.81 | 6.82±0.72 | 5.01±1.55 | 11.21±0.35 | ABC transporter ATP-binding protein |
| *Q3H53_000242* | 3.23±1.02 | 6.62±1.08 | 6.85±0.6 | 5.74±2.56 | 3.93±0.23 | type II toxin-antitoxin system HicA family toxin |
| *Q3H53_000244* | 0±0 | 0.24±0.07 | 0.67±0.02 | 0.24±0.23 | 0.58±0.12 | TdeIII family type II restriction endonuclease |
| *Q3H53_000361* | 13.39±0.47 | 33.1±4.36 | 31.59±0.62 | 29.17±7.36 | 54.38±2.29 | type II toxin-antitoxin system HicA family toxin |
| *Q3H53_000379* | 5.67±0.97 | 11.48±0.43 | 12.79±0.62 | 11.45±1.53 | 13.76±0.12 | glycosyltransferase family 1 protein |
| *Q3H53_000380* | 1.24±0.54 | 3.22±0.77 | 3.04±0.22 | 2.65±0.87 | 4.9±0.23 | glycosyltransferase family 1 protein |
| *Q3H53_000442* | 4.6±0.48 | 10.18±1.74 | 12.29±0.42 | 10.98±3.5 | 16.88±0.21 | restriction endonuclease |
| *Q3H53_000487* | 5.09±0.43 | 13.06±2.22 | 12.24±0.5 | 8.05±1.8 | 8.13±0.51 | metalloregulator ArsR/SmtB family transcription factor |
| *Q3H53_000492* | 11.6±1.94 | 27.96±4.1 | 27.07±0.01 | 36.47±7.11 | 40.82±0.98 | DUF2203 domain-containing protein |
| *Q3H53_000565* | 0.6±0.12 | 3.71±0.68 | 3.78±0.22 | 2.54±1.28 | 4.09±0.19 | hypothetical protein |
| *Q3H53_000605* | 0±0 | 2.31±0.04 | 1.58±0.09 | 1.25±0.5 | 0.99±0.01 | hypothetical protein |

| GeneID | 0.05:B | 0.05:A | 0.20:B | 0.20:A | 0.05:E | Product |
|---|---|---|---|---|---|---|
| *Q3H53_000610* | 32.72±3.39 | 76.7±6.93 | 79±5.25 | 48.89±10.28 | 86.32±2.42 | hypothetical protein |
| *Q3H53_000631* | 9.04±0.37 | 24.06±7.87 | 21.75±0.89 | 17.16±3.33 | 24.07±0.82 | hypothetical protein |
| *Q3H53_000697* | 1.51±0.97 | 4.42±2.1 | 5.08±0.13 | 2.27±0.33 | 3.81±0.15 | hypothetical protein |
| *Q3H53_000752* | 10.18±1.21 | 22.15±1.48 | 35.69±0.09 | 17.05±3.16 | 29.49±3.36 | hypothetical protein |
| ***Q3H53_000913*** | **2.53±0.34** | **5.18±0.84** | **5.44±0.45** | **4.31±0.87** | **4.23±0.48** | **DNA replication/repair protein RecF** |
| *Q3H53_000966* | 0±0 | 1.96±1.22 | 2.74±0.16 | 1.24±0.67 | 3.58±0.31 | hypothetical protein |
| *Q3H53_000988* | 1.33±0.18 | 2.97±0.75 | 2.78±0.05 | 3.41±1.73 | 3.54±0.55 | hypothetical protein |
| *Q3H53_000992* | 0±0 | 0.81±0.18 | 0.52±0.04 | 0.33±0.21 | 0.2±0 | DALR anticodon-binding domain-containing protein |
| ***Q3H53_001148*** | **7.43±5.07** | **2.6±0.02** | **3.18±0.08** | **2.4±0.62** | **1.46±0.1** | **nitrate ABC transporter permease** |
| *Q3H53_001195* | 2.47±0.15 | 7.84±0.6 | 6.15±0.25 | 5.89±2.63 | 8.74±0.16 | DUF29 domain-containing protein |
| *Q3H53_001206* | 3.29±0.63 | 9.22±1.46 | 6.94±0.27 | 4.41±0.57 | 4.53±0.32 | hypothetical protein |
| *Q3H53_001258* | 0.35±0.25 | 1.17±0.11 | 1.17±0.24 | 0.49±0.2 | 0.73±0.05 | DUF87 domain-containing protein |
| *Q3H53_001295* | 7.76±3.41 | 22.5±7.65 | 26.01±1.79 | 17.67±2.06 | 32.03±1.64 | transcriptional regulator |
| *Q3H53_001326* | 2.46±0.97 | 9.89±0.4 | 7.87±0.28 | 4.67±0.03 | 3.13±0.39 | hypothetical protein |
| *Q3H53_001329* | 3.32±0.24 | 7.41±0.82 | 7.45±0.22 | 7.58±1.42 | 7.46±0.12 | metallophosphoesterase family protein |
| *Q3H53_001348* | 9.4±1.21 | 25.32±3.49 | 24.47±2.95 | 21.34±4.41 | 22.84±0.46 | hypothetical protein |
| *Q3H53_001496* | 8.01±0.49 | 20.33±1.62 | 19.79±0.07 | 17.66±3.44 | 25.3±1.3 | response regulator transcription factor |
| *Q3H53_001516* | 0.33±0.47 | 1.49±0.45 | 1.72±0.27 | 0.32±0.13 | 0.96±0.1 | hypothetical protein |

| GeneID | 0.05:B | 0.05:A | 0.20:B | 0.20:A | 0.05:E | Product |
|---|---|---|---|---|---|---|
| *Q3H53_001562* | 1.23±1.11 | 4.88±1.36 | 3.25±0.47 | 3.18±1.76 | 2.24±0.4 | DUF433 domain-containing protein |
| *Q3H53_001609* | 2.74±1.86 | 7.75±2.83 | 9.52±0.19 | 7±2.32 | 13.71±0.24 | type II toxin-antitoxin system VapC family toxin |
| *Q3H53_001610* | 10.29±2.74 | 28.15±0.89 | 25.4±0.14 | 21.76±6.96 | 35.75±0.64 | hypothetical protein |
| *Q3H53_001615* | 0±0 | 0.74±0.14 | 0.78±0.08 | 0.08±0.12 | 0.36±0 | endonuclease domain-containing protein |
| *Q3H53_001678* | 2.38±0.53 | 6.2±0.18 | 8.52±0.16 | 8.27±2.12 | 7.97±0.2 | GAF domain-containing protein |
| *Q3H53_001730* | 0.25±0.35 | 1.23±0.49 | 1.92±0.11 | 0.85±0.11 | 0.91±0.07 | ABC transporter ATP-binding protein |
| *Q3H53_001760* | 60.67±36.67 | 18.81±1.43 | 21.31±0.99 | 32.24±3.27 | 14.79±0.5 | hydrogenase maturation nickel metallochaperone HypA |
| *Q3H53_001761* | 64.23±34.66 | 26.13±0.23 | 26.68±0.13 | 38.14±2.28 | 25.66±0.23 | agmatinase family protein |
| *Q3H53_001797* | 16.29±6.61 | 36.77±3.04 | 35.11±0.19 | 27.29±8.42 | 43.4±0.96 | transposase |
| *Q3H53_001826* | 11.18±2.18 | 23.22±3.24 | 22.45±1.12 | 16.32±3.59 | 22.98±1.45 | hypothetical protein |
| *Q3H53_001969* | 0.64±0.9 | 3.17±0.46 | 2.97±0.67 | 2.2±0.28 | 3.33±0.54 | type II toxin-antitoxin system VapC family toxin |
| *Q3H53_001970* | 7.83±1.06 | 17.12±2.55 | 21.67±1.9 | 18.1±5.53 | 20.52±0.58 | DUF2281 domain-containing protein |
| *Q3H53_002000* | 6.36±2.87 | 20.07±1.07 | 22.88±0.04 | 17.14±7.23 | 34.06±0.96 | type II toxin-antitoxin system VapC family toxin |
| ***Q3H53_002006*** | **2.15±0.41** | **6.78±2.69** | **4.88±0.32** | **2.57±0.89** | **3.94±0.23** | **6-carboxytetrahydropterin synthase QueD** |
| *Q3H53_002046* | 0.6±0.33 | 1.6±0.16 | 2.89±0.1 | 0.86±0.08 | 1.25±0.05 | hypothetical protein |
| *Q3H53_002117* | 0±0 | 1.01±0.85 | 0.92±0.07 | 0.74±0.16 | 0.96±0.12 | hypothetical protein |
| *Q3H53_002119* | 2.46±1.56 | 8.46±4.07 | 6.76±0.67 | 6.53±0.23 | 6.78±0.09 | hypothetical protein |
| *Q3H53_002146* | 22.35±2.71 | 46.37±7.05 | 62.12±1.38 | 44.36±4.33 | 71.28±0.18 | DUF29 domain-containing protein |

| GeneID | 0.05:B | 0.05:A | 0.20:B | 0.20:A | 0.05:E | Product |
|---|---|---|---|---|---|---|
| *Q3H53_002155* | 0±0 | 0.42±0.13 | 0.82±0.05 | 0.13±0.18 | 0.36±0.3 | PIN domain-containing protein |
| *Q3H53_002159* | 1.45±0.42 | 9.98±0.6 | 12.05±0.56 | 8.22±4.09 | 22.97±1.53 | restriction endonuclease |
| *Q3H53_002160* | 25.72±13.14 | 79.33±7.9 | 68.01±2.66 | 56.94±19.91 | 74.33±1.4 | helix-turn-helix transcriptional regulator |
| *Q3H53_002221* | 0.81±0.1 | 1.87±0.08 | 2.06±0.1 | 1.12±0.34 | 1.58±0.06 | SAM-dependent methyltransferase |
| *Q3H53_002282* | 0±0 | 0.35±0.1 | 0.85±0.19 | 0.55±0.32 | 1.07±0.01 | hypothetical protein |
| *Q3H53_002286* | 0.24±0.34 | 1.03±0.06 | 1.25±0.07 | 0.51±0.13 | 0.88±0.19 | hypothetical protein |
| *Q3H53_002299* | 16.47±1.7 | 34.67±4.69 | 39.96±1.26 | 23.34±6.63 | 41.35±0.8 | pyridoxal phosphate-dependent aminotransferase |
| *Q3H53_002305* | 0.23±0.32 | 0.66±0.07 | 2.7±0.02 | 0.23±0.02 | 1.09±0.13 | hypothetical protein |
| *Q3H53_002360* | 6.59±0.39 | 3.16±1.87 | 2.93±0.78 | 4.41±0.16 | 2.54±0.05 | cadmium resistance transporter |
| *Q3H53_002381* | 2.33±0.8 | 4.91±1.67 | 4.81±0.07 | 2.65±0.17 | 3.58±0.14 | hypothetical protein |
| *Q3H53_002424* | 20.86±3.34 | 56.14±5.56 | 50.24±0.35 | 49.28±11.77 | 55.57±2.73 | type II toxin-antitoxin system RelE/ParE family toxin |
| *Q3H53_002425* | 19.57±5.11 | 47.87±2.29 | 61.05±2.22 | 42.52±2.03 | 59.76±2.93 | HigA family addiction module antitoxin |
| *Q3H53_002471* | 0.6±0.33 | 1.6±0.16 | 2.89±0.1 | 0.86±0.08 | 1.25±0.05 | hypothetical protein |
| *Q3H53_002481* | 5.99±0.17 | 14.34±1.79 | 13.35±1.05 | 10.99±2.02 | 13.76±0.29 | hypothetical protein |
| *Q3H53_002539* | 14.82±2.92 | 40.47±8.05 | 32.1±0.08 | 17.75±4.39 | 30.71±0.79 | AI-2E family transporter |
| *Q3H53_002673* | 1.7±0.58 | 5.78±0.26 | 4.04±0.62 | 5.73±2.17 | 2.9±0.68 | calcium-binding protein |
| *Q3H53_002858* | 2.43±1.49 | 6±1.39 | 5.99±0.2 | 4.47±0.16 | 4.04±0.03 | hypothetical protein |
| *Q3H53_002894* | 3.01±0.25 | 6.7±0.83 | 8.54±0.28 | 8.9±1.52 | 11.08±0.25 | tetratricopeptide repeat protein |

| GeneID | 0.05:B | 0.05:A | 0.20:B | 0.20:A | 0.05:E | Product |
|---|---|---|---|---|---|---|
| *Q3H53_003139* | 7.65±3.54 | 29.59±6.86 | 15.56±0.15 | 11.53±2.53 | 11.58±0.08 | hypothetical protein |
| *Q3H53_003156* | 0.14±0.2 | 1.27±0.71 | 1.99±0 | 0.99±0.13 | 1.74±0.36 | fluoride efflux transporter CrcB |
| *Q3H53_003212* | 0±0 | 1.09±0.2 | 0.63±0.14 | 2.16±0 | 1.34±0.76 | hypothetical protein |
| *Q3H53_003215* | 1.72±1.79 | 5.23±0.34 | 5.15±0.95 | 3.25±0.24 | 4.09±0.16 | hypothetical protein |
| *Q3H53_003390* | 676.38±156.19 | 1354.68±80.65 | 2092.79±84.12 | 1615.53±205.25 | 2784.27±56.24 | DUF4327 family protein |
| *Q3H53_003391* | 248.28±15.24 | 521.23±31.25 | 785.93±26.01 | 620.36±38.09 | 1112.66±26.8 | efflux RND transporter periplasmic adaptor subunit |
| *Q3H53_003437* | 2.39±1.71 | 7.95±4.45 | 6.76±0.75 | 5.02±2.46 | 5.21±0.7 | hypothetical protein |
| *Q3H53_003458* | 1.93±0.8 | 3.87±0.37 | 3.98±0.18 | 3.97±2.4 | 5.3±0.21 | polysaccharide biosynthesis C-terminal domain-containing protein |
| *Q3H53_003460* | 1.31±0.33 | 2.65±0.25 | 3.14±0.14 | 2.83±0.8 | 4±0.06 | glycosyltransferase family 1 protein |
| *Q3H53_003483* | 0.82±0.34 | 2.58±1.17 | 1.97±0.02 | 1.61±0.12 | 1.65±0.04 | glycosyltransferase family A protein |
| *Q3H53_003566* | 0.87±1.23 | 2.19±0.68 | 2.42±0.04 | 1.72±0.22 | 1.73±0.19 | type II toxin-antitoxin system VapC family toxin |
| *Q3H53_003585* | 6.69±0.75 | 17±0.14 | 19.05±0.31 | 15.52±3.03 | 25.24±0.61 | hypothetical protein |
| *Q3H53_003586* | 13.29±3.34 | 28.13±2.76 | 31.22±0.87 | 20.76±1.22 | 35.18±0.57 | DUF4926 domain-containing protein |
| *Q3H53_003587* | 7.91±0.74 | 20.68±1.3 | 24.97±0.2 | 13.35±0.74 | 28.74±1.05 | hypothetical protein |
| *Q3H53_003590* | 5.05±1.43 | 10.54±0.36 | 10.58±0.16 | 7.23±1.29 | 12.72±1.03 | hypothetical protein |
| *Q3H53_003622* | 1.52±0.29 | 5.36±0.94 | 4.74±0.59 | 3.26±0.07 | 4.59±0.75 | DUF3146 family protein |
| *Q3H53_003660* | 1.82±0.25 | 6.21±1.52 | 5.02±0.35 | 3.7±1.68 | 6.76±0.27 | AAA family ATPase |
| *Q3H53_003672* | 2.25±1.57 | 4.71±0.32 | 6.61±0.51 | 5±1.62 | 8.49±0.64 | hypothetical protein |

| GeneID | 0.05:B | 0.05:A | 0.20:B | 0.20:A | 0.05:E | Product |
|---|---|---|---|---|---|---|
| *Q3H53_003680* | 1.16±0.34 | 2.83±0.62 | 2.36±0.09 | 1.33±0.59 | 1.52±0.15 | lysylphosphatidylglycerol synthase transmembrane domain-containing protein |
| *Q3H53_003740* | 26.81±1.9 | 71.68±7.71 | 68.37±0.57 | 61.63±12.13 | 91.39±0.31 | YggS family pyridoxal phosphate-dependent enzyme |
| *Q3H53_003766* | 9.79±0.7 | 23.69±0.2 | 25.81±0.03 | 21.84±4.9 | 18.87±0.48 | hypothetical protein |
| *Q3H53_003829* | 1.58±0.61 | 3.77±0.98 | 4.15±0.09 | 2.28±0.17 | 3.11±0.02 | element excision factor XisI family protein |
| *Q3H53_003830* | 2.17±1 | 4.8±1.48 | 0.61±0.04 | 2.19±1.75 | 0.85±0.3 | element excision factor XisH family protein |
| *Q3H53_003831* | 0.9±0.17 | 2.63±1.45 | 2.82±0.16 | 2.2±0.52 | 2.27±0.34 | hypothetical protein |
| *Q3H53_003877* | 0±0 | 1.5±0.4 | 2.84±0.01 | 0.65±0.26 | 1.03±0.36 | hypothetical protein |
| *Q3H53_003958* | 0±0 | 1.76±0.24 | 2.39±0.31 | 1.27±0.51 | 1.25±0.13 | hypothetical protein |
| *Q3H53_004004* | 5.06±0.05 | 11.13±1.12 | 11.85±0.06 | 7.39±1.69 | 7.26±1 | Mov34/MPN/PAD-1 family protein |
| *Q3H53_004008* | 3.3±0.98 | 9.43±2.67 | 9.4±0.13 | 4.9±1.16 | 6.8±0.05 | hypothetical protein |
| *Q3H53_004065* | 0.27±0.39 | 1.91±0.05 | 2.37±0.22 | 2.05±0.74 | 5.02±0.41 | HNH endonuclease |
| *Q3H53_004109* | 1.1±0.07 | 2.89±0.87 | 2.58±0.36 | 2.37±0.13 | 2.89±0.09 | nitroreductase family protein |
| ***Q3H53_004127*** | **10.58±0.11** | **22.58±2.82** | **24.1±0.13** | **15.02±3.77** | **20.78±0.47** | **DNA repair protein RecO** |
| *Q3H53_004352* | 11.42±6.6 | 4.44±1.53 | 5.66±0.45 | 9.49±0.09 | 6.12±0.36 | hypothetical protein |
| *Q3H53_004406* | 1.29±0.06 | 3.09±1.15 | 4.05±0.01 | 2.75±1.65 | 2.64±0.14 | hypothetical protein |
| *Q3H53_004426* | 10.52±1.92 | 22.31±0.41 | 23.11±0.01 | 19.74±4.71 | 30.75±0.91 | tRNA (adenosine(37)-N6)-dimethylallyltransferase MiaA |
| *Q3H53_004730* | 3.61±0.45 | 7.77±1.65 | 8.81±0.31 | 11.1±0.76 | 13.21±0.58 | hypothetical protein |
| *Q3H53_004866* | 0.6±0.33 | 1.6±0.16 | 2.89±0.1 | 0.86±0.08 | 1.25±0.05 | hypothetical protein |

**Table S4.** Enriched KEGG pathways. nDEGs represent the number of differentially expressed genes in this KEGG pathway. BgGs represent the number of annotated genes in this KEGG pathway. Padj represents Benjamini-Hochberg adjusted *p* value.

| **Comparison** | **Up/down** | **Layer1** | **Layer2** | **Layer3** | **KO** | **nDEGs** | **BgGs** | ***p*** | **Padj** | **ID of DEGs** |
|---|---|---|---|---|---|---|---|---|---|---|
| 0.05:A/B | Up | Genetic Information Processing | Replication and repair | Homologous recombination | ko03440 | 2 | 19 | 0.040455 | 0.606819 | Q3H53_000913\|Q3H53_004127 |
| | | Metabolism | Metabolism of cofactors and vitamins | Folate biosynthesis | ko00790 | 2 | 21 | 0.04865 | 0.364878 | Q3H53_002006\|Q3H53_002069 |
| | Down | Environmental Information Processing | Membrane transport | ABC transporters | ko02010 | 5 | 75 | 0.001116 | 0.005579 | Q3H53_000107\|Q3H53_000108\|Q3H53_001148\|Q3H53_001149\|Q3H53_004461 |
| | | Metabolism | Energy metabolism | Nitrogen metabolism | ko00910 | 2 | 16 | 0.015455 | 0.038637 | Q3H53_000107\|Q3H53_000108 |
| 0.20:A/B | Up | Cellular Processes | Cellular community - prokaryotes | Quorum sensing | ko02024 | 2 | 33 | 0.026881 | 0.161286 | Q3H53_000568\|Q3H53_002062 |
| | Down | Metabolism | Energy metabolism | Sulfur metabolism | ko00920 | 3 | 17 | 0.001784 | 0.026758 | Q3H53_001536\|Q3H53_001537\|Q3H53_001538 |

| Comparison | Up/down | Layer1 | Layer2 | Layer3 | KO | nDEGs | BgGs | *p* | Padj | ID of DEGs |
|---|---|---|---|---|---|---|---|---|---|---|
| B:0.20/0.05 | Up | Genetic Information Processing | Replication and repair | Homologous recombination | ko03440 | 2 | 19 | 0.038053 | 0.456639 | Q3H53_000913\|Q3H53_004127 |
| | Down | Metabolism | Energy metabolism | Nitrogen metabolism | ko00910 | 2 | 16 | 0.008167 | 0.040833 | Q3H53_000108\|Q3H53_003417 |
| 0.05:E/A | Up | Cellular Processes | Cellular community - prokaryotes | Quorum sensing | ko02024 | 3 | 33 | 0.056776 | 0.999832 | Q3H53_000568\|Q3H53_000948 \|Q3H53_002062 |
| | Down | NULL | NULL | NULL | NULL | NULL | NULL | NULL | NULL | NULL |
